# Divalent cation depletion enhances neuronal excitability through CaSR-dependent modulation of threshold channels

**DOI:** 10.64898/2025.12.07.692819

**Authors:** Konstantina Mylonaki, Matías Alvarez-Saavedra, Michaël Russier, Dominique Debanne, Salvatore Incontro

## Abstract

External calcium ([Ca^²⁺^]□) and magnesium ([Mg^²⁺^]□) concentrations fluctuate across physiological and pathological brain states. For example, [Ca^²⁺^]□ decreases during intense neuronal activity and epilepsy, whereas it rises during sleep. Similarly, [Mg^²⁺^]□ varies with the sleep/wake cycle and is reduced in epilepsy. Lowering either [Ca^2+^]_e_ or [Mg^2+^]_e_ increases intrinsic excitability and hyperpolarizes the action potential (AP) threshold, yet the underlying mechanisms remain unclear. Here, we confirm that reducing [Ca^2+^]_e_ or [Mg^2+^]_e_ enhances intrinsic excitability and hyperpolarizes the AP threshold of CA1 pyramidal neurons. Physiological reductions in [Mg^²⁺^]□ (0.8 → 0.4 mM) have minimal effect, whereas decreases from supraphysiological levels (2.0 → 0.4 mM) robustly increase excitability. Using pharmacology and CRISPR/Cas9 gene editing, we identify the calcium-sensing receptor (CaSR) as a key mediator of these effects. The calcilytic NPS-2143 mimics and largely occludes both the intrinsic excitability increase and the AP-threshold hyperpolarization, while genetic reduction of CaSR produces similar outcomes. We further show that AP-threshold hyperpolarization induced by low divalent cations involves both Kv1 and Nav1.2 channels. Together, these findings reveal CaSR as a critical link between external divalent cation levels and intrinsic neuronal excitability through the modulation of Kv1 and Nav channels.

## Introduction

External calcium concentration ([Ca^2+^]_e_) fluctuates between 1.2 and 0.8 mM across the sleep/wake cycle (Ding *et al*., 2016) and can drop to 0.1 mM during intense neural activity such as that observed during epilepsy. Physiological calcium levels deeply modify synaptic plasticity rules induced by spike-timing (Inglebert *et al*., 2020; Chindemi *et al*., 2022). The effects of both physiological and pathological reduction in [Ca^2+^]_e_ have been studied in the context of neuronal excitability (Lu *et al*., 2010), particularly regarding their impact on the action potential (AP) threshold (Segal, 2018). A decrease in [Ca^2+^]_e_ elevates intrinsic neuronal excitability and hyperpolarizes the AP threshold, but the underlying mechanisms remain a topic of debate. One proposed mechanism involves the lesser activation of SK channels (Segal, 2018). However, SK channels typically require calcium influx triggered by an AP for activation and their effects are generally absent on the first AP (Sourdet *et al*., 2003). Moreover, internal calcium buffering with BAPTA does not suppress the hyperpolarization of the AP threshold (Forsberg *et al*., 2025), suggesting that alternative pathways may be involved.

External magnesium concentration [Mg^2+^]_e_ also varies during the sleep/wake cycle from ∼0.70 mM during wakefulness to ∼0.83 mM during sleep, reaching up to 1.14 mM during anesthesia (Ding *et al*., 2016). External magnesium plays fundamental role in brain physiology as it blocks NMDA receptors at hyperpolarized potentials (Ascher & Nowak, 2009) and facilitates GABA_A_ receptor activity (Möykkynen *et al*., 2001). Intracellular magnesium is also an essential co-factor for adenosine triphosphate (ATP) production (Yamanaka *et al*., 2016) and for gap junction plasticity (Palacios-Prado *et al*., 2013, 2014). In neurons, levels of [Mg^2+^]_e_ are tightly regulated by Mg^2+^-permeant channels and transporters, including transient receptor potential melastatin-like 7 (TRPM7), cylin M 3 (CNNM3), and magnesium transporter 1 (MagT1) located at the plasma membrane (de Baaij *et al*., 2015). Neuronal activity increases intracellular Mg^2+^ concentration (Yamanaka *et al*., 2015) while reduced [Mg^2+^]_e_ levels are reported in epileptic patients (Kirkland *et al*., 2018). Raising [Mg^2+^]_e_ has been shown to elevate AP threshold and reduce intrinsic excitability (Frankenhaeuser & Meves, 1958; Dribben *et al*., 2010). However, the mechanisms by which [Mg^2+^]_e_ controls neuronal excitability are still poorly understood.

The calcium sensing receptor (CaSR) is a G-protein coupled receptor (GPCR) that detects fluctuations in both [Ca^2+^]_e_ and [Mg^2+^]_e_, and is expressed in neurons (Ruat & Traiffort, 2013; Giudice *et al*., 2019). In addition to divalent ions such as Ca^2+^ and Mg^2+^, CaSR is also activated by polyamines such as spermine (Quinn *et al*., 1997) and L-amino acid (Conigrave *et al*., 2000). Furthermore, CaSR is activated by the calcimimetic cinacalcet and inhibited by the calcilytic NPS-2143 (Saidak *et al*., 2009). CaSR has been shown to inhibit both glutamate release (Phillips *et al*., 2008) and intrinsic neuronal excitability (Lu *et al*., 2010; Mattheisen *et al*., 2018). Importantly, CaSR activity can be inhibited by a drop in [Ca^2+^]_e_ (Martiszus *et al*., 2021). However, pharmacological studies targeting CaSR have yielded inconsistent results: both calcilytics like NPS-2143 and calcimimetics like cinacalcet were reported to inhibit the voltage-gated sodium current in dissociated neurons (Mattheisen *et al*., 2018). Thus, the precise mechanism by which CaSR modulates the AP threshold and neuronal excitability remain unresolved.

In this study, we revisited this question using CA1 pyramidal neurons from organotypic slice cultures of rat hippocampus. Unlike findings in dissociated cultures, we show that CaSR inhibition via the calcilytic NPS-2143 increases neuronal excitability by hyperpolarization the AP threshold. NPS-2143 mimics and partially occludes the excitability enhancement induced by low [Ca^2+^]_e_ and [Mg^2+^]_e_. In contrast, the calcimimetic cinacalcet reduces neuronal excitability primarily through a decrease in input resistance. Additionally, inhibition of Kv1 channels with dendrotoxin-I partially blocks the AP threshold hyperpolarization while pharmacological blockade or genetic reduction of Nav1.2 channels strongly reduces the increase in excitability. Taken together these results suggest that reduction in [Ca^2+^]_e_ and [Mg^2+^]_e_ enhance neuronal excitability via a CaSR-dependent modulation of Kv1 and Nav channels.

## Results

### Lowering [Ca^2+^]_e_ and [Mg^2+^]_e_ enhances intrinsic neuronal excitability

Whole-cell recordings from CA1 pyramidal neurons were obtained in organotypic slice cultures of rat hippocampus after 8-15 days *in vitro*. To examine the effects of extracellular divalent ions on neuronal excitability, extracellular calcium and magnesium concentrations ([Ca^²⁺^]□ and [Mg^²⁺^]□) were reduced from 3.0/2.0 to 0.6/0.4 mM. Neuronal excitability was tested during this transition using both constant current pulses and current steps of increasing amplitude to assess the input–output relationship (**Figure 1A**; **Figure S1**).

**Figure 1.**
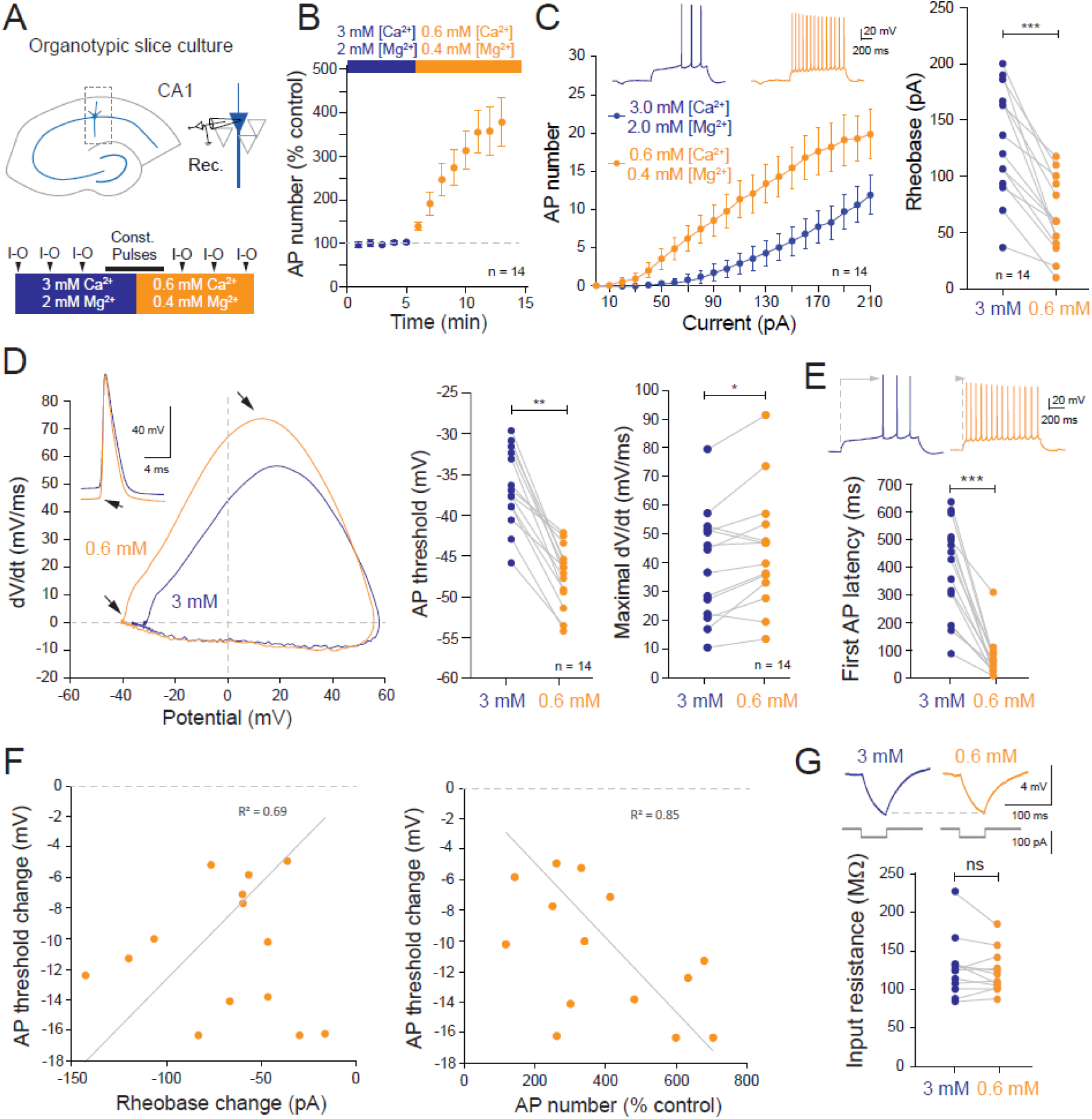
Lowering [Ca^2+^]_e_ and [Mg^2+^]_e_ enhances intrinsic neuronal excitability in CA1 pyramidal neurons. A. Experimental design. B. Time course of the increased excitability induced by lowering [Ca^2+^]_e_ and [Mg^2+^]_e_. C. Comparison of input-output curves in 3.0 / 2.0 mM [Ca^2+^]_e_ and [Mg^2+^]_e_ with 0.6 / 0.4 mM. D. Hyperpolarization of the AP threshold and increase in the rising phase of the AP in the presence of low divalent ions. E. Reduction of the first spike latency. F. Correlations of AP threshold and rheobase changes (linear regression, y = 0.126x, R^2^=0.69) and, AP threshold and AP number changes (linear regression, y =-0.024x, R^2^=0.85). G. No change in input resistance were observed following lowering [Ca^2+^]_e_ and [Mg^2+^]_e_. *, Wilcoxon test, p<0.05; **, Wilcoxon test p<0.01, ***, Wilcoxon test p<0.001.

On average, the transition from high to low divalent cation concentrations increased excitability; the number of action potentials (APs) evoked by a constant current increased nearly fourfold (**Figure 1B**). The input-output curve was shifted leftward due to a reduction in rheobase (from 131 ± 13 to 63 ± 9 pA, n = 14; **Figure 1C**). The AP threshold was markedly hyperpolarized (from-36.7 ± 1.3 to-47.6 ± 1.0 mV, n = 14; **Figure 1D**), and the maximal rising slope of the AP was enhanced (from 38.8 ± 5.1 to 44.3 ± 5.5 mV/ms, n = 14; **Figure 1D**). The increase in excitability was associated with a substantial reduction in first-spike latency (from 404 ± 45 to 76 ± 20 ms; **Figure 1E**). Notably, larger hyperpolarization of the AP threshold was associated with greater reduction in rheobase and stronger excitability increase (**Figure 1F**). No significant change in input resistance was observed following the transition from high divalent to low divalent cation concentrations (from 125.3 ± 9.5 to 120.6 ± 6.9 MΩ, n = 14; **Figure 1G**).

To determine whether calcium reduction alone accounts for this effect, we selectively lowered [Ca^²⁺^]□ while maintaining [Mg^²⁺^]□ constant. Lowering [Ca^2+^]_e_ increased intrinsic neuronal excitability to ∼160% of control levels (**Figure 2A**). The rheobase decrease was comparable to that observed when both ions were reduced (-43 ± 9 pA, n = 13 vs.-68 ± 9 pA, n = 14; **Figure 2B**), but the AP threshold hyperpolarization was smaller (-4.6 ± 0.7 mV, n = 13 vs.-10.8 ± 1.1 mV in 0.6/0.4 mM, n = 14; **Figure 2C**). Despite this, the degree of threshold hyperpolarization remained correlated with both rheobase reduction and the increase in AP number (**Figure 2D**).

**Figure 2.**
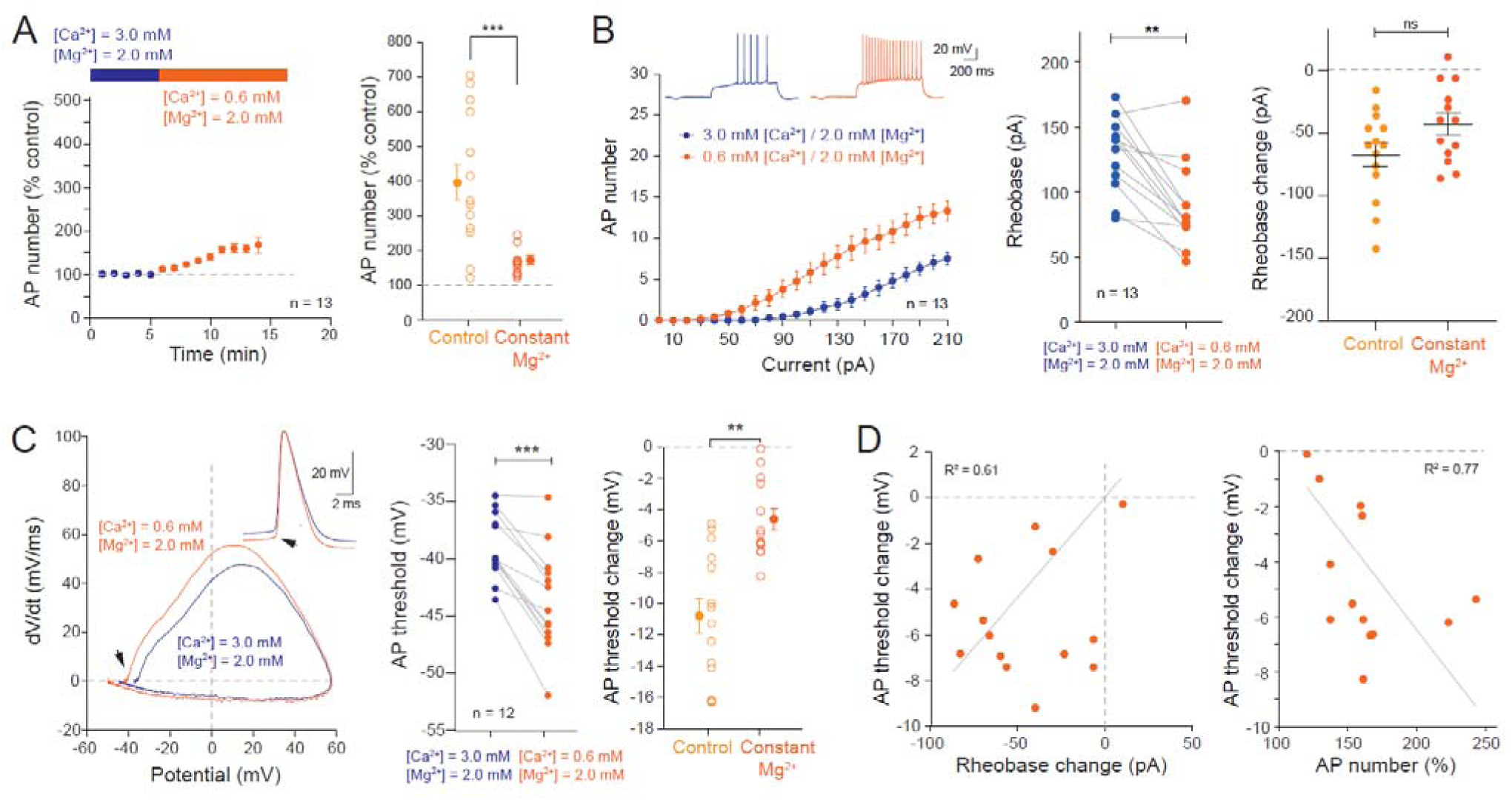
In constant [Mg^2+^]_e_, lowering [Ca^2+^]_e_ from 3.0 to 0.6 mM induces smaller changes in excitability. A. Left, time-course of the increased excitability during the transition from 3.0 mM to 0.6 mM [Ca^2+^]_e_ in constant extracellular magnesium ([Mg^2+^] = 2.0 mM). The increase in excitability is significantly lower in constant magnesium (***, Mann-Whitney test, p<0.001). B. Left, input-output curves in 3.0 mM [Ca^2+^]_e_ and 2.0 magnesium (blue dots) and in 0.6 mM calcium and 2.0 magnesium (orange dots). Middle, rheobase change induced low calcium alone; **, Wilcoxon test, p<0.01. Right, comparison of the rheobase change in control and in constant magnesium; ns, Mann-Whitney U-test, p>0.1. C. Left, phase plots of the first AP in 3.0 mM / 2.0 mM (blue) and in 0.6 mM / 2.0 mM (orange). Middle, AP threshold changes; ***, Wilcoxon test, p<0.001. Right, comparison of the AP threshold changes in control (i.e., change in both [Ca^2+^] and [Mg^2+^] versus in constant [Mg^2+^]. D. Correlation of the AP threshold change with rheobase change (left; y = 0.078x, R² = 0.61) and AP number (right; y =-0.065x, R²=0.77).

As the physiological [Ca^2+^]_e_ in the brain is ∼1.3 mM (Jones & Keep, 1988; Silver & Erecińska, 1990), we next tested whether similar effects occur when [Ca^²⁺^]□ and [Mg^²⁺^]□ are reduced from 1.3/0.8 mM to 0.6/0.4 mM. This reduction enhanced excitability to ∼200% of the control (**Figure 3A** & **3B**), accompanied by a decrease in the rheobase (from 68 ± 6 pA to 34 ± 4 pA, n = 16; **Figure 3C**) and a modest hyperpolarization of AP threshold ( ∼3 mV: from-48.3 ± 0.8 to-51.8 ± 0.8 mV, n = 16; **Figure 3D**). Again, AP threshold shifts correlated with the rheobase reduction and excitability changes (**Figure 3E**).

**Figure 3.**
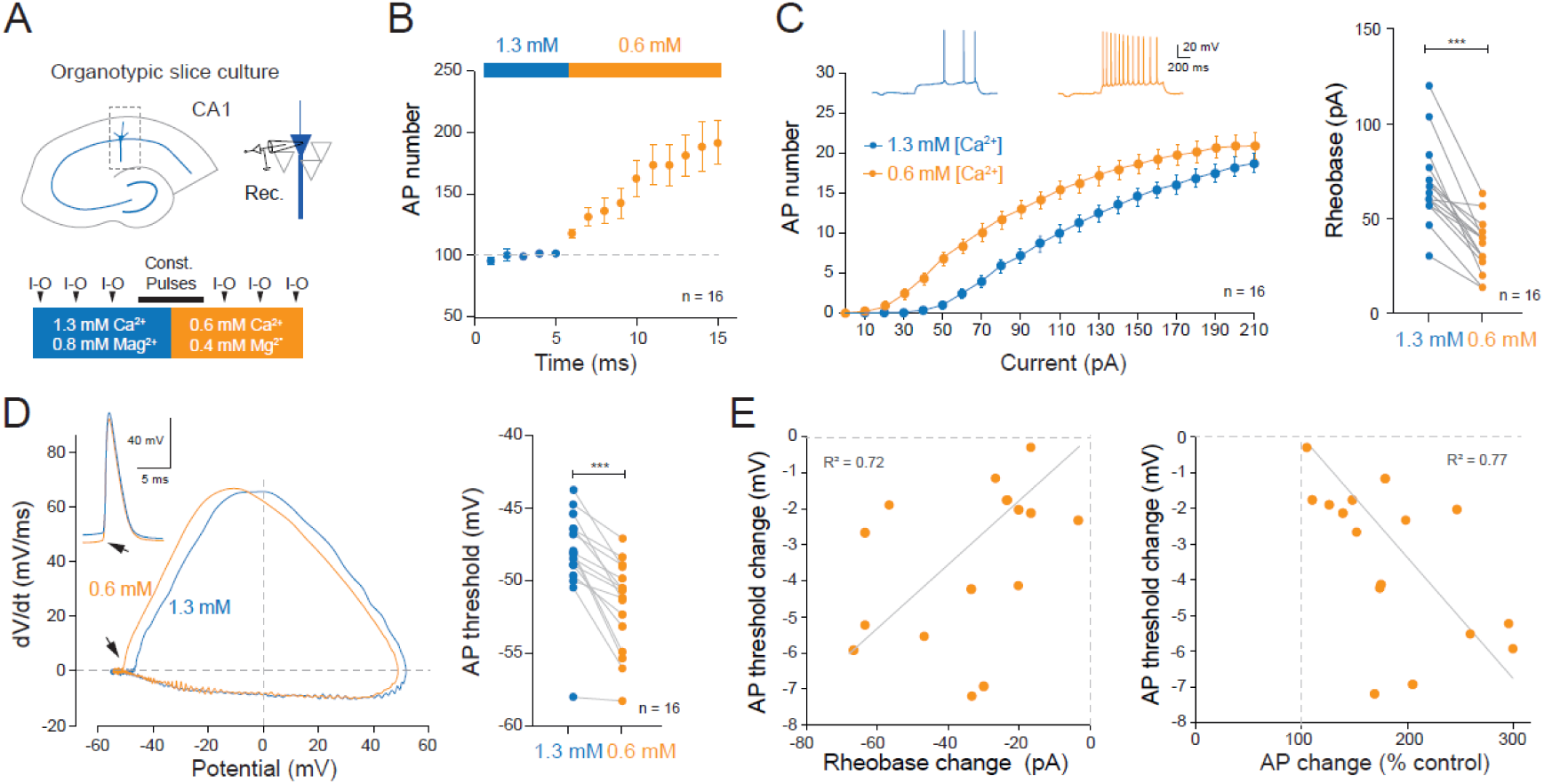
Lowering of [Ca^2+^]_e_ and [Mg^2+^]_e_ in the physiological range enhances intrinsic neuronal excitability. A. Experimental design and protocol. B. Time course of the increased excitability induced by lowering [Ca^2+^]_e_ (from 1.3 to 0.6 mM) and [Mg^2+^]_e_ (from 0.8 to 0.4 mM). C. Comparison of input-output curves from 1.3 mM [Ca^2+^]_e_ / 0.8 mM [Mg^2+^]_e_ magnesium to 0.6 [Ca^2+^]_e_ / 0.4 mM [Mg^2+^]_e_. D. Hyperpolarization of the AP threshold. Left, phase plots of the first AP in 1.3 mM [Ca^2+^]_e_ / 0.8 mM [Mg^2+^]_e_ and in 0.6 mM [Ca^2+^]_e_ / 0.4 mM [Mg^2+^]_e_. E. Correlation of the AP threshold change with rheobase change (left) and AP change (right). ***, Wilcoxon test, p<0.001.

When [Ca^²⁺^]□ alone was reduced from 1.3 to 0.6 mM (with [Mg^²⁺^]□ kept at 0.8 mM), excitability increased similarly (∼200% of the control; from 4.5 ± 0.3 to 8.2 ± 0.8 APs, *n* = 14; **Figure 4A**). The rheobase decrease (−32 ± 7 pA, *n* = 14) and threshold hyperpolarization (−2.6 ± 0.3 mV, *n* = 14) were comparable to those observed when both ions were reduced (**Figure 4B–C**). These results indicate that, at physiological levels, magnesium plays a minor role in modulating excitability changes induced by reduced divalent cation concentrations (**Figure 4D**).

**Figure 4.**
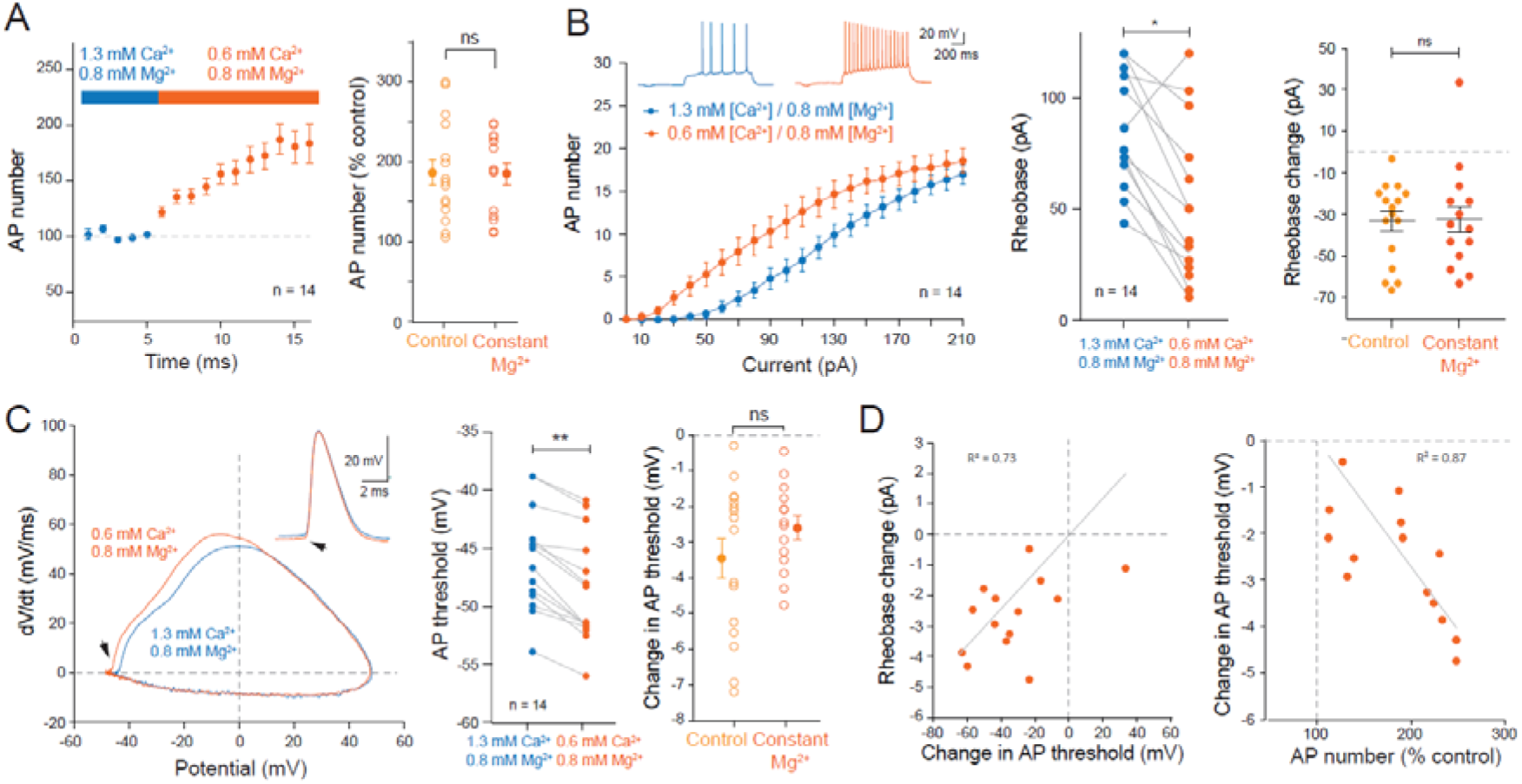
In constant [Mg^2+^]_e_, lowering [Ca^2+^]_e_ from 1.3 mM to 0.6 mM increases excitability similarly. A. Left, time-course of the effect of lowering [Ca^2+^]_e_ from 1.3 mM to 0.6 mM in the presence of constant [Mg^2+^]_e_. Right, comparison of the excitability changes in control vs. in constant [Mg^2+^]_e_. ns, Mann-Whitney, p>0.1. B. Left, input-output curve before and after lowering [Ca^2+^]_e_ from 1.3 to 0.6 mM. Middle, change in the rheobase induced by low [Ca^2+^]_e_. Right, comparison of the rheobase reduction in control condition (i.e., reduction of both [Ca^2+^]_e_ and [Mg^2+^]_e_) vs. in constant [Mg^2+^]_e_. *, Wilcoxon test, p<0.05. ns, Mann-Whitney U-test, p>0.1. C. Analysis of the spike threshold. Left, phase plots of the AP before and after the reduction in [Ca^2+^]_e_. Middle, hyperpolarization of the AP threshold. Right, comparison of the AP threshold changes in control and in constant [Mg^2+^]_e_.

When comparing high (3.0/2.0 mM) versus physiological (1.3/0.8 mM) divalent cation concentrations, the AP threshold was ∼10 mV more hyperpolarized (−47.2 ± 0.7 mV, *n* = 30 vs. −37.9 ± 0.8 mV, *n* = 27; Mann–Whitney U test, *p* < 0.001; **Figure S2**) and the rheobase was significantly reduced (75.6 ± 4.4 pA vs. 132.2 ± 7.9 pA; **Figure S2**). The plot of the rheobase as a function of the AP threshold for each cell confirmed a joint reduction of both parameters in physiological [Ca^2+^]_e_ and [Mg^2+^]_e_ (**Figure S2**).

Together, these results demonstrate that reductions in external divalent cations reliably enhance intrinsic excitability primarily through coordinated shifts in rheobase and AP threshold.

### CaSR inhibition by NPS-2143 enhances excitability and occludes low-divalent cation-mediated effects

CaSR expression was confirmed in CA1 pyramidal neurons (**Figure S3**) and has been suggested to be involved in the regulation of intrinsic excitability. To assess its contribution to intrinsic excitability, we applied the calcilytic NPS-2143 (10 µM). NPS-2143 increased AP firing to ∼130% of control (**Figure 5A**), shifted the input–output curve slightly leftward, and decreased the rheobase (from 117 ± 6 to 97 ± 7 pA, n = 13; **Figure 5B**). Importantly, the AP threshold was hyperpolarized (from-40.5 ± 0.7 to-41.5 ± 0.7 mV, n = 13; **Figure 5C**) without any change in input resistance (from 160 ± 10 to 168 ± 8 MΩ, n = 13; **Figure 5D**). These results indicate that NPS mimics the effects of reduced [Ca^2+^] _e._

**Figure 5.**
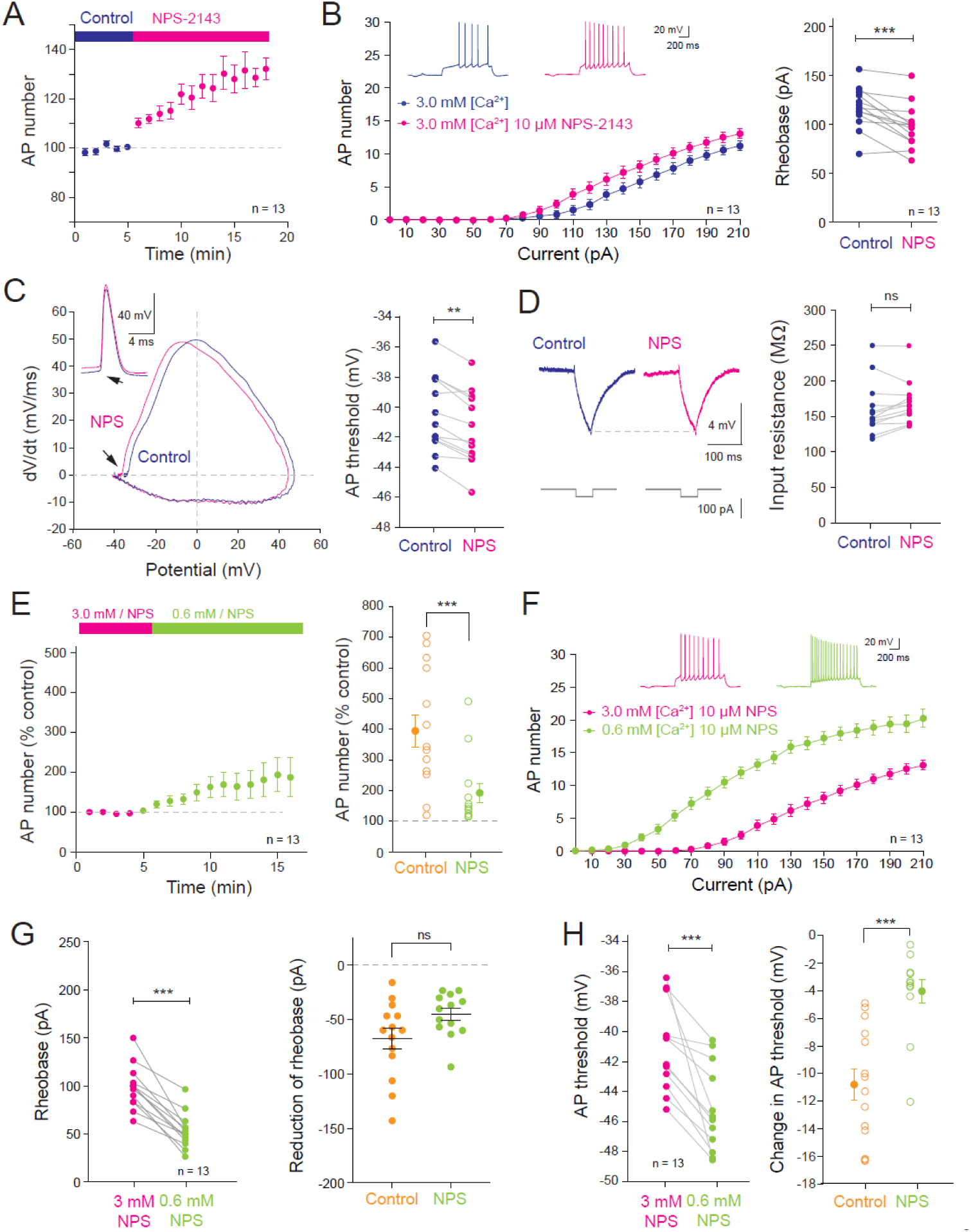
NPS-2143, mimics and partially occludes the effects of lowering [Ca^2+^] and [Mg^2+^]. A. Left, time course of the excitability change induced by NPS-2143 in high calcium and magnesium ([Ca^2+^]_e_ = 3.0 mM and [Mg^2+^]_e_ = 2.0 mM). B. Left, input-output curves before and after bath application of NPS-2143. Right, change in rheobase. ***, Wilcoxon test, p<0.001. C. Analysis of AP threshold. Left, phase-plots of the first AP before and after 15 minutes of NPS-2143 application. D. No change in input resistance observed. ns, Wilcoxon test, p>0.05. E. Left, enhanced excitability induced by low calcium in the presence of NPS-2143. Right, comparison of the excitability increases in control and NPS-2143. ***, Mann-Whitney U-test, p<0.001. F. Input-output curves and rheobase in 3.0 mM and 0.6 mM calcium in the presence NPS-2143. G. Left, rheobase reduction in the presence of NPS-2143. Right, comparison of the rheobase change. ***, Wilcoxon test, p<0.001. ns, Mann-Whitney U-test, p>0.1. H. Left, hyperpolarization of the AP threshold in the presence of NPS-2143. ***, Wilcoxon test, p<0.001. Right, comparison of the AP threshold hyperpolarization in control and in NPS-2143. ***, Mann-Whitney, p<0.001.

Next, we tested whether NPS-2143 occludes the effect of low calcium and magnesium on neuronal excitability. In the presence of NPS-2143, the increase in excitability was markedly smaller than in control (191 ± 31%, n = 13 vs. 395 ± 52% in control, n = 14; **Figure 5E**). The input-output curve shifted leftward (**Figure 5F**) and the rheobase decreased (from 97 ± 7 pA to 55 ± 5 pA, n = 13; **Figure 5G**), but the magnitude of rheobase change did not significantly differ from control (**Figure 5G**). The AP threshold hyperpolarization was, however, strongly reduced in the presence of NPS-2143, (-4.0 ± 0.8 mV, n = 13 vs.-10.8 ± 1.1 mV, n = 14 in control; **Figure 5H**) indicating that CaSR inhibition largely occludes the low-calcium effect on excitability.

When applied under physiological [Ca^²⁺^]□/[Mg^²⁺^]□, NPS-2143 produced a smaller effect on excitability (∼140% of control; **Figure S4**) compared to high [Ca^²⁺^]□/[Mg^²⁺^]□ conditions. In these physiological conditions, AP threshold and rheobase were not significantly altered (**Figures S4**).

Finally, when [Ca^²⁺^]□/[Mg^²⁺^]□ were reduced from 1.3/0.8 mM to 0.6/0.4 mM in the presence of NPS-2143, AP firing still increased (**Figure S4**), but the rheobase reduction was significantly smaller than in control (−20 ± 3 pA vs. −34 ± 5 pA; **Figure S4**), and AP threshold remained unchanged (**Figure S4**).

Collectively, these findings indicate that NPS-2143 partially occludes the increase in excitability induced by reduced extracellular calcium and magnesium.

### Effects of cinacalcet on the enhanced excitability induced by low divalent cations

We next examined whether the calcimimetic cinacalcet, a CaSR agonist, can attenuate the increase in intrinsic excitability induced by lowering external divalent cation concentrations. Bath application of cinacalcet (20 µM) reduces neuronal excitability of CA1 pyramidal cells through the reduction in input resistance and not through the depolarization of the AP threshold (**Figure S5**). Specifically, cinacalcet decreased input resistance by ∼10% (i.e., from 143 ± 14 to 128 ± 12 MΩ, n = 7; **Figure S5**).

In the presence of cinacalcet, the elevation in AP number during the transition from 3.0/2/0 to 0.6/0.4 mM [Ca^²⁺^]□ / [Mg^²⁺^]□was significantly reduced compared to control conditions (250 ± 38%, n = 12 vs. 395 ± 52% in control n = 14; **Figure 6A**), although input resistance remained constant throughout the transition (**Figure 6B**). The input-output curve was shifted leftward (**Figure 6C**) but neither the reduction in rheobase (**Figure 6D**) nor the AP threshold hyperpolarization (**Figure 6E**) differed significantly from control conditions.

**Figure 6.**
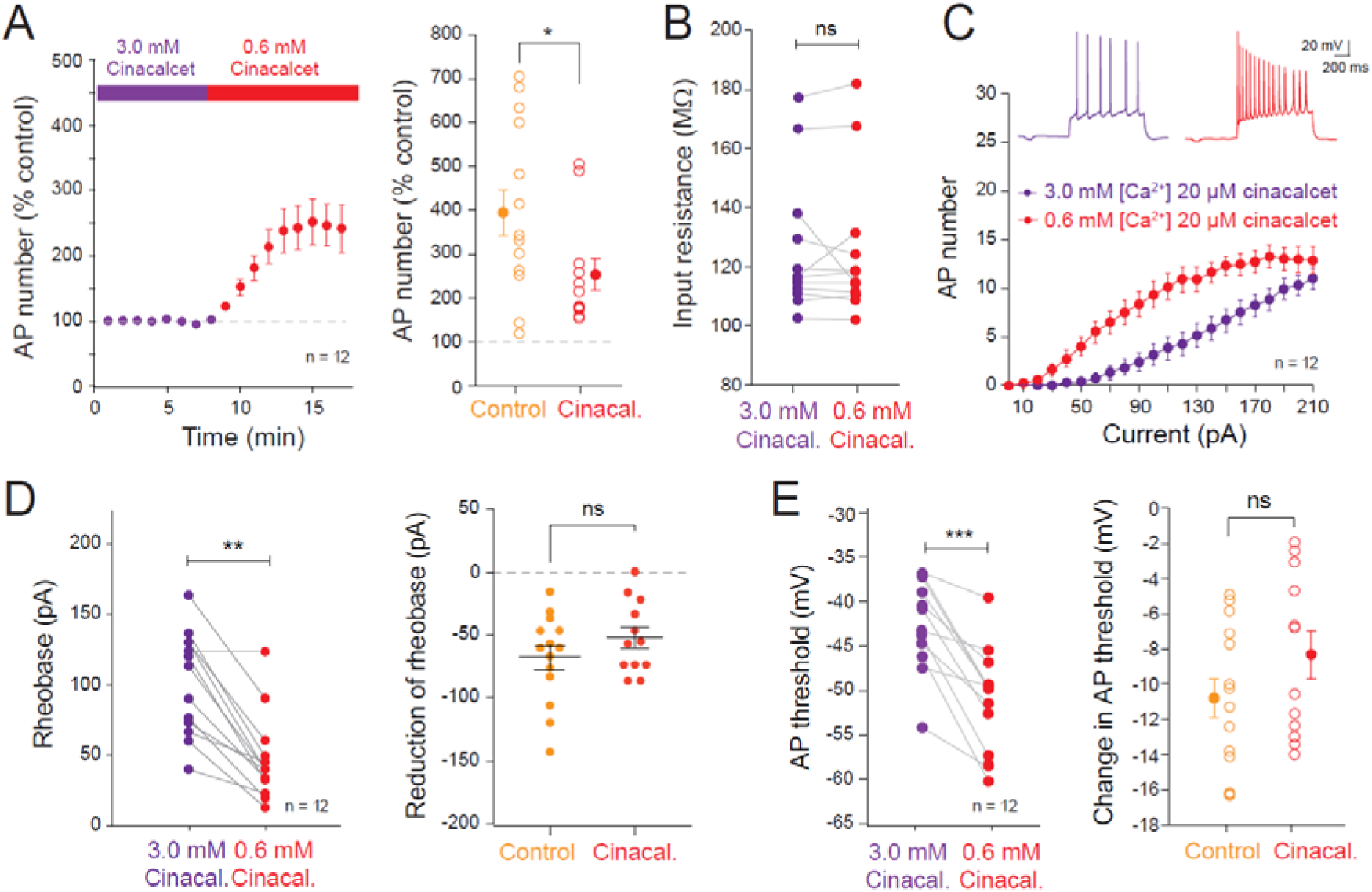
Cinacalcet reduces the effects of low divalent cations on intrinsic excitability. A. Left, time-course of the elevation in intrinsic excitability induced by lowering calcium and magnesium in the presence of 20 µM cinacalcet. Right, comparison of the AP number in control and in cinacalcet. *, Mann-Whitney U-test, p<0.05. B. No change in input resistance is observed. C. Input-output curves in the presence of the cinacalcet. D. Left, rheobase change in the presence of cinacalcet. **, Wilcoxon test, p<0.01. Right, comparison of the rheobase change in control and cinacalcet. ns, Mann-Whitney U-test, p>0.1. E. Left, change in AP threshold induced by the reduction of external calcium and magnesium in the presence of cincalcet. ***, Wilcoxon test, p<0.001. Right, comparison of the AP threshold changes in control and in cinacalcet. ns, Mann-Whitney U-test, p>0.1.

We next tested the effect of cinacalcet in physiological [Ca^2+^]_e_ and [Mg^2+^]_e_. In the presence of cinacalcet, the reduction in the rheobase was found to be significantly reduced but not the AP threshold (**Figure S6**).

Taken together, these results indicate that cinacalcet partially occludes the enhancement of intrinsic excitability caused by reduction in external [Ca^2+^]_e_ and [Mg^2+^]_e_.

### Genetic impairment of CaSR partially occludes the AP threshold changes

We next examined whether the increase in intrinsic excitability induced by low [Ca^²⁺^]□ and [Mg^²⁺^]□ is altered by genetically impairing CaSR function. To down-regulate CaSR expression, we used CRISPR/Cas9 (Jinek *et al*., 2012; Cong *et al*., 2013) and electroporated CA1 pyramidal neurons with plasmids encoding CRISPR-CaSR together with EGFP (**Figure 7A**). The efficiency of the CRISPR was tested on dissociated neurons transfected either with an empty plasmid (control) or with the CRISPR-CaSR plasmid. Western blot analysis revealed a clear reduction in CaSR protein levels in CRISPR-CaSR–transfected neurons (**Figure S7**), confirming effective gene knockdown.

**Figure 7.**
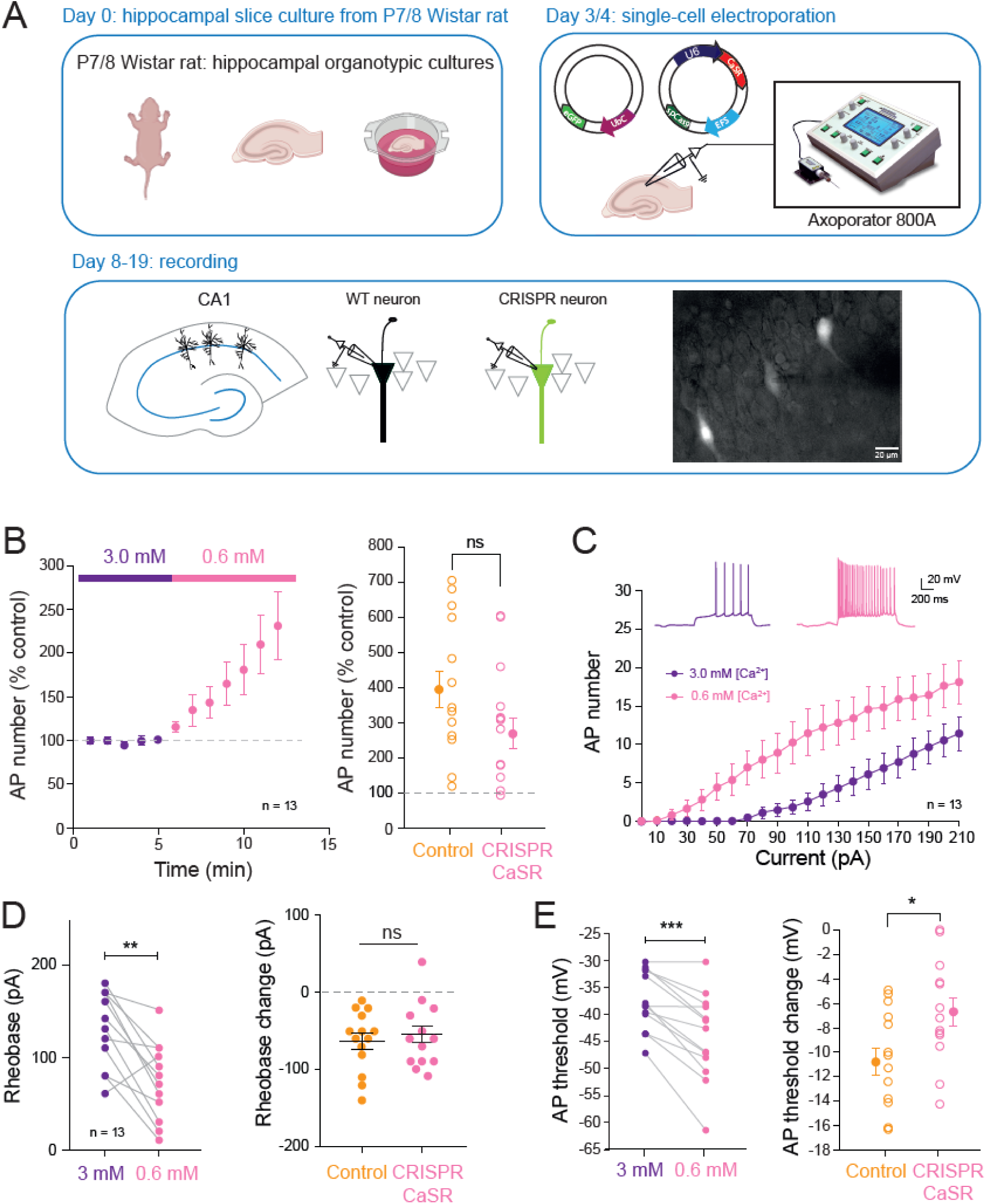
Genetic impairment of CaSR reduces the AP hyperpolarization. A. Protocol. B. Left, time-course of the increased excitability in neurons transfected with CRISPR CaSR. Right, comparison of the AP number in control and in CRISPR CaSR neurons. ns, Mann-Whitney U-test, p>0.1. C. Input-output curves in CRISPR CaSR-transfected neurons. D. Left, rheobase change. **, Wilcoxon test, p<0.01. Right, comparison of the rheobase changes in control and in CRISPR CaSR transfected neurons. ns, Mann-Whitney U-test, p>0.1. E. Left, AP threshold changes. ***, Wilcoxon test, p<0.001. Right comparison of the AP threshold changes in control and in CRISPR CaSR neurons. * Mann-Whitney U-test, p<0.05.

In CRISPR-CaSR transfected neurons, the transition from 3.0/2.0 mM to 0.6/0.4 mM [Ca^²⁺^]□ / [Mg^²⁺^]□ elevated the AP number to a level comparable to the control (**Figure 7B**) and the rheobase change was not significantly reduced (-57 ± 12 pA, n = 12 vs.-62 ± 10 pA, n = 14; **Figure 7C** and **7D**). However, the hyperpolarization of the AP threshold was significantly reduced (-6.7 ± 1.2, n = 12 vs.-10.8 ± 1.1 m in control, n = 14; **Figure 7E**). In physiological [Ca^2+^]_e_ and [Mg^2+^]_e_, the reduction of divalent ions in CRISPR-CaSR transfected neurons led to an elevation of excitability that was like the control (**Figure S8**). Both the rheobase and AP threshold changes were not significantly different from control. Taken together, these results strongly support a major role of CaSR in the excitability increase during reduction of [Ca^2+^]_e_.

### Blocking Kv1 channels partly occludes the AP threshold changes

The reduction in the first AP latency generally indicates that Kv1 channels are downregulated (Campanac *et al*., 2013; Duménieu *et al*., 2025). To test whether Kv1 channels contribute to the effects of lowered divalent cations, we repeated the initial experiment in the presence of dendrotoxin-I (DTx-I), a selective blocker of Kv1.1, Kv1.2, and Kv1.6 subunits. In the presence of DTx-I, lowering [Ca^2+^]_e_ and [Mg^2+^]_e_ to 0.6/0.4 mM increased neuronal excitability, but to a similar extent compared to control conditions (**Figure 8A**). Rheobase decreased by 60 pA in DTx-I vs. 76 pA, in control conditions (**Figure 8B** and **Figure 8C**). First-spike latency was similarly reduced in the presence of DTx-I compared to control (**Figure 8D**). However, AP threshold hyperpolarization was significantly attenuated in the presence of DTx-I compared with control (-6.8 ± 1.3 mV n = 12 in DTx-I vs.-10.8 ± 1.1 mV in control, n = 14; **Figure 8E**).

**Figure 8.**
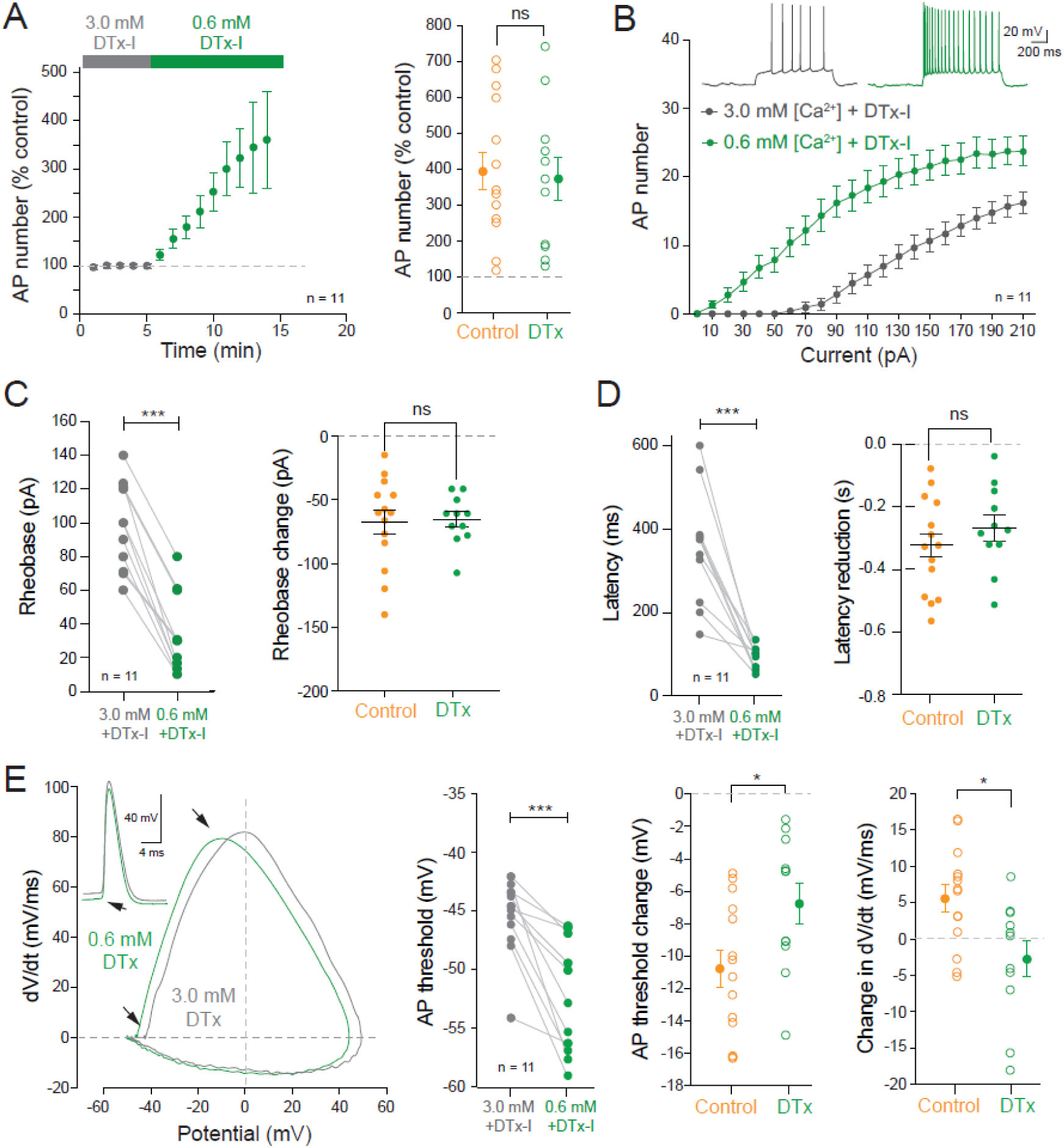
Inhibition of Kv1 channels with DTx-I partially occludes the hyperpolarization of the AP threshold induced by low calcium. A. Transition from 3.0/2.0 mM to 0.6/0.4 mM in the presence of 100 nM of the Kv1 channel blocker, DTx-I. Left, time-course of the enhanced excitability induced by low divalent cations. Right, comparison with control. ns, Mann-Whitney U-test, p>0.1. B. Input-output curves. C. Analysis of rheobase. Left, reduction of the rheobase induced by low divalent cations. ***, Wilcoxon test, p<0.001. Right, comparison with control. ns, Mann-Whitney U-test, p>0.1. D. Analyses of the first AP latency (left) and comparison with control (right). ***, Wilcoxon, p<0.001. ns, Mann-Whitney U-test p>0.1. E. Analyses of AP phase plot and AP threshold. Left, phase plots of APs in 3.0/2.0 mM and 0.6/0.4 mM in the presence of DTx-I. Middle left, hyperpolarization of AP threshold. ***, ***, Wilcoxon, p<0.001. Middle right, change in AP threshold compared with control. *, Mann-Whitney, p<0.05. Right, change in maximal dV/dt. *, Mann-Whitney, p<0.05.

These results indicate that reduced Kv1 channel function contributes, at least in part, to the increased excitability and AP threshold hyperpolarization induced by low [Ca^2+^]_e_ and [Mg^2+^]_e_.

### Freezing Nav1.2 partially occludes the AP threshold changes

As Kv1 channels are not responsible for the total hyperpolarization of the AP threshold induced by lowering [Ca^2+^]_e_ and [Mg^2+^]_e_, we next examined the contribution of Nav1.2 and Nav1.6 channels using selective pharmacological blockers. Application of Huwentoxin IV (HWTx IV), a Nav1.2-selective channel blocker (Minassian *et al*., 2013; Filipis *et al*., 2023) reduced the Nav current by ∼15% in 3.0/2.0 mM (**Figure 9A**), without altering the input-output curve and the rheobase (**Figure 9B**). HWTx-IV reduced the maximal speed of the depolarizing phase of the AP (**Figure 9C**). In the presence of HWTx-IV, the drop of [Ca^2+^]_e_ and [Mg^2+^]_e_ enhanced the excitability and reduced the rheobase comparable to the control (**Figure 9D** & **9E**) but significantly reduced the hyperpolarization of the AP threshold (**Figure 9F**). HWTx-IV produced comparable effects at 1.3/0.8 mM, partially occluding the rheobase reduction while leaving the AP threshold hyperpolarization largely unchanged (**Figure S9**). Together, these results indicate that Nav1.2 contributes to the excitability increase triggered by reduced extracellular divalent cations.

**Figure 9.**
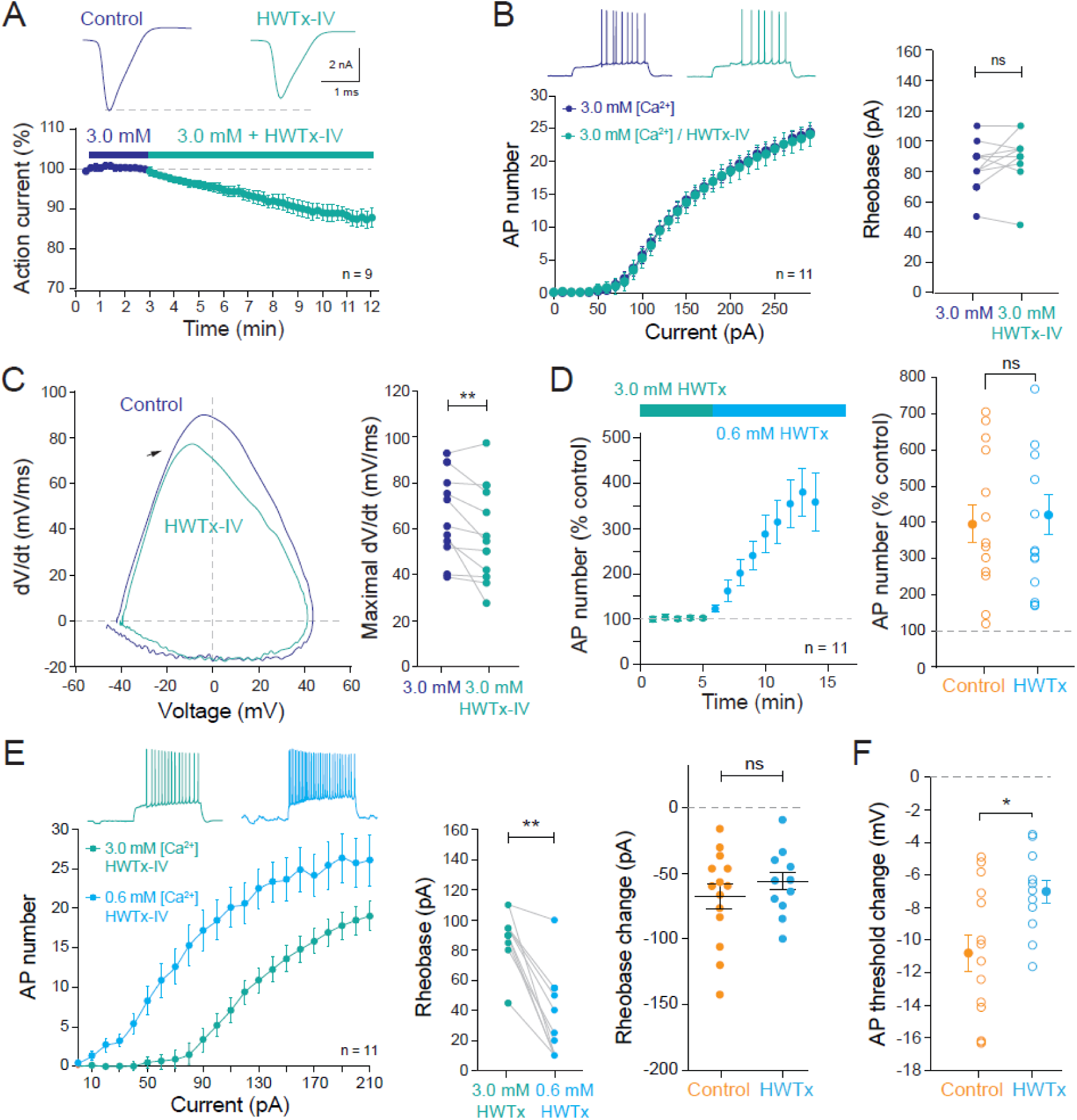
Nav1.2 channel blockade reduces the hyperpolarization of the AP threshold. A. Reduction of the Nav current by 300 nM of HWTx-IV, a Nav1.2-selective toxine. Top, representative currents evoked by a depolarization from-80 to 0 mV in a CA1 pyramidal neuron. Bottom, time-course of the reduction. B. HWTx-IV does not alter the input output curve. Left, input-output curves. Right rheobase. ns, Wilcoxon test, p>0.1. C. AP phase-plot (left) and maximal dV/dt changes (right) induced by low divalent cations in the presence of HWTx-IV. **, Wilcoxon test, p<0.01. D. Time course of the enhanced excitability induced by low divalent cations (left) and comparison with control (right). ns, Mann-Whitney U-test, p>0.1. E. Left, input-output curves in high and low divalent cations. Middle, changes in the rheobase. **, Wilcoxon test, p<0.01. Right, comparison of the AP threshold hyperpolarization in HWTx and in control. *, Mann-Whitney U-test, p <0.05.

We next tested the role of Nav1.6 using 4,9-anhydro-tetrodotoxin (ATTx), a Nav1.6-selective blocker (Hargus *et al*., 2013; Li *et al*., 2023). ATTx reduced the Nav current amplitude by ∼20% in 3.0/2.0 mM (**Figure 10A**) and decreased the maximal dv/dt of the AP (**Figure 10B** & **10C**), but did not significantly affect rheobase, excitability, or AP threshold (**Figure 10 D-F**). These observations suggest that Nav1.6 does not substantially contribute to the divalent cation-dependent increase in excitability.

**Figure 10.**
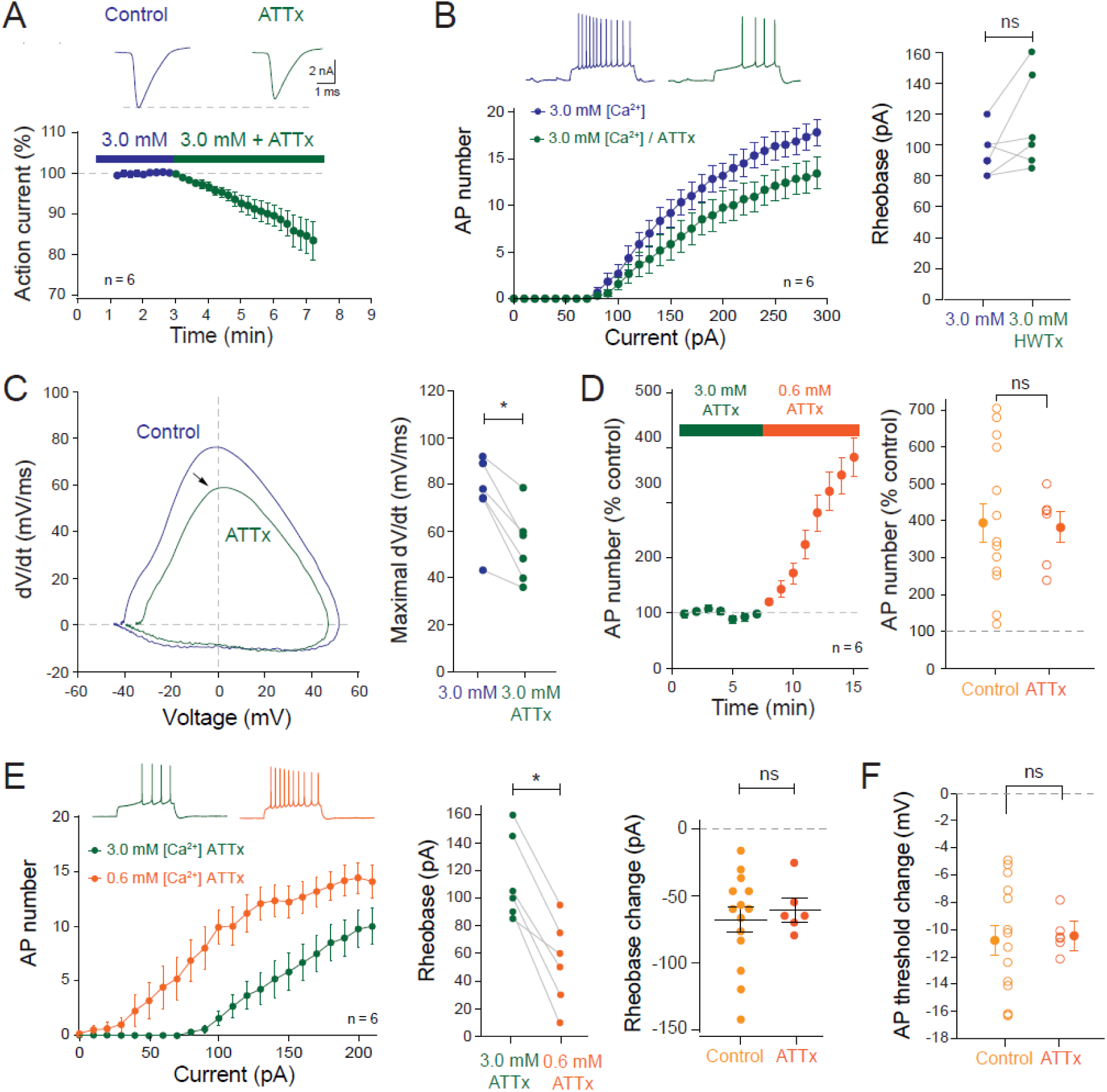
The Nav1.6 channel blocker, ATTx doies not occlude the AP therhold change. A. ATTx-induced reduction of the sodium current evoked by a step of current from - 80 to 0 mV. B. Change in input-output curves induced by ATTx (left) and rheobase analysis (right). ns, Wilcoxon test, p>0.1. C. AP phase-plots (left) and maximal dV/dt analysis (right). *, Wilcoxon test, p<0.05. D. Time course of the excitability enhancement induced by low divalent cations (left) and comparision of the AP number in control and in the presence of ATTx (right). ns, Mann-Whitney U-test, p>0.1. E. Left, input-output curves in high and low divalent cations. Middle left, rheobase analysis. *, Wilcoxon test, p<0.05. Middle right, comparison of the rheobase changes in control and ATTx. ns, Mann-Whitney U-test, p>0.1. F. Hyperpolarization of AP threshold in control and in ATTx. ns, Mann-Whitney U-test, p>0.1.

To further test this idea, we genetically impaired Nav1.1/1.2 and Nav1.6 using CRISPR/Cas9. CA1 neurons in which Nav1.1/Nav1.2 were deleted displayed a significant reduction in the AP threshold hyperpolarization induced by lowering [Ca^²⁺^]□ and [Mg^²⁺^]□ (**Figure 11**), confirming a key role for Nav1.2 in this process. In contrast, CRISPR-mediated reduction of Nav1.6 expression had no detectable effect on the number of APs, the rheobase change, or the AP threshold shift following divalent cation reduction (**Figure S10**). Finally, under physiological [Ca^²⁺^]□ and [Mg^²⁺^]□, Nav1.1/Nav1.2 CRISPR neurons also exhibited a significantly reduced rheobase change (**Figure S11**), reinforcing the central contribution of Nav1.2 to the excitability increase induced by low extracellular calcium. Together, these findings identify Nav1.2 as a central mediator of the divalent cation-dependent shift in AP threshold, providing a mechanistic link between extracellular calcium sensing receptors and intrinsic excitability.

**Figure 11.**
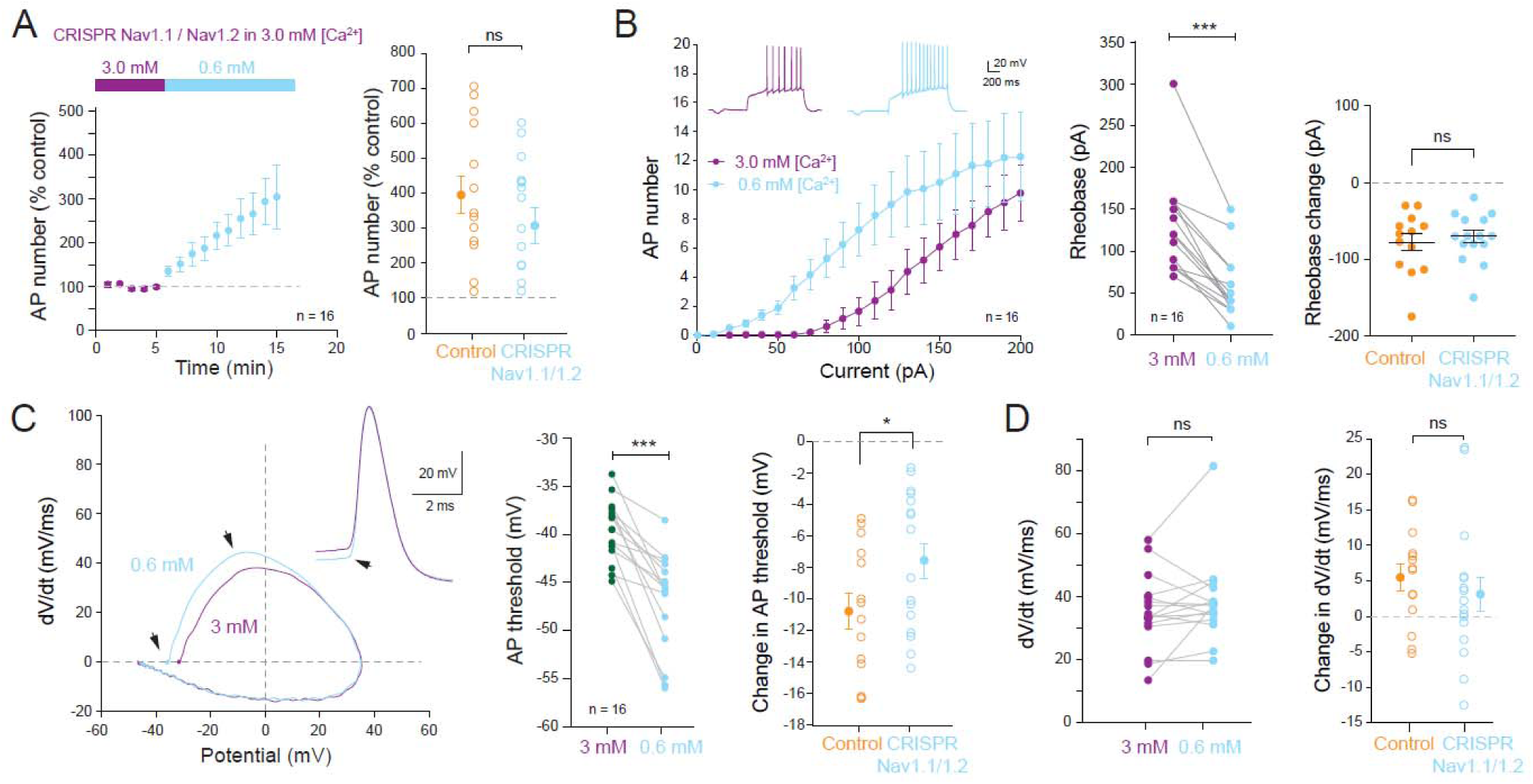
Genetic impairment of Nav1.1/1.2 reduces AP threshold hyperpolarization. A. Time-course of the excitability enhancement induced by low divalent cations in CRISPR Nav1.1/Nav1.2 neurons. Input-output curve and rheobase analysis of neurons electroporated with CRISPR targeting Nav1.1/1.2. B. Transition from 3.0 to 0. 6 mM. C. AP phase-plots (left) and AP threshold analysis (middle and right). Compared to control the AP threshold hyperpolarization is significantly reduced in CRISPR Nav1.1/1.2. ***, Wilcoxon test, p<0.001. **, Mann-Whitney U-test, p<0.05. D. Analysis of the maximal dV/dt. Left, change in maximal dV/dt during the transition from high to low divalent cations. ns, Wilcoxon test, p>0.1. Right, change in dV/dt compared to control. ns, Mann-Whitney U-test, p>0.1.

## Discussion

We report here that reducing extracellular calcium and magnesium markedly enhances neuronal excitability in CA1 pyramidal neurons by hyperpolarizing the AP threshold. In contrast to previous work in dissociated cultures (Mattheisen *et al*., 2018), inhibition of CaSR with the calcilytic NPS-2143 increases neuronal excitability and largely occludes the effect of low divalent cation concentrations. Conversely, the calcimimetic cinacalcet suppresses neuronal excitability, not by modulating Nav channels as previously suggested (Lindner *et al*., 2022), but primarily by decreasing input resistance. Finally, blocking Kv1 channels with dendrotoxin-I, inhibiting Nav1.2 channels with HWTx or genetically reducing Nav1.1/1.2 partially occludes the hyperpolarizing shift in AP threshold induced by reduced [Ca^2+^]_e_ and [Mg^2+^]_e_, whereas Nav1.6 blockade or knockdown has no significant effect. Together, these findings define a CaSR–Nav1.2–Kv1 axis that controls intrinsic excitability during fluctuations in extracellular divalent cation concentrations.

### Low [Ca^2+^]_e_ and [Mg^2+^]_e_ increase excitability in CA1 hippocampal neurons

Historically, surface changes screening has been proposed as the primary explanation for the increased intrinsic excitability observed during low [Ca^2+^]_e_ (Gilbert & Ehrenstein, 1969). Because negatively charged outer membrane phospholipids bind divalent cations, reducing [Ca^²⁺^]□ (and especially [Mg^²⁺^]□) results in substantial relief of surface charge screening. In our experiments, the drop from 3.0/2.0 mM to 0.6/0.4 mM total divalent cation concentration represents a fivefold reduction and is therefore expected to produce a large shift in AP threshold. This reduction is proportionally smaller when starting from 1.3/0.8 mM, where the total divalent cation concentration decreases from 2.1 mM to 1.0 mM. When [Mg^2+^]_e_ is kept constant, the total divalent cation concentration drops from 5 mM to 2.6 mM (i.e., a difference of 2.4 mM) in one case and from 2.1 mM to 1.4 mM (i.e., 0.7 mM) in the other case. Despite these differences, rheobase consistently dropped by ∼50–60% and the AP threshold hyperpolarized by ∼5 mV, indicating that surface charge effects alone cannot account for the full magnitude of the excitability increase.

Intracellular calcium has also been implicated in threshold regulation through activation of SK channels (Segal, 2018). However, a recent study showed that chelating intracellular Ca^²⁺^ with BAPTA does not prevent threshold hyperpolarization (Forsberg *et al*., 2025).

### CaSR plays a central role in the augmentation of intrinsic excitability

In both high and low [Ca^²⁺^]□, the calcilytic NPS-2143 mimicked the effect of low divalent cations, increasing AP number by ∼30–40% and hyperpolarizing the AP threshold by ∼0.4–1.0 mV. Moreover, NPS-2143 partially occluded the excitability increase caused by reducing [Ca^²⁺^]□ and [Mg^²⁺^]□ from 3.0/2.0 mM to 0.6/0.4 mM.. This occlusion was weaker when starting from 1.3/0.8 mM, consistent with a smaller CaSR activation range. NPS-2143 significantly attenuated rheobase changes but did not alter the threshold shift itself.

Genetic reduction of CaSR expression similarly diminished the AP threshold hyperpolarization at 3.0/2.0 mM but not at 1.3/0.8 mM, further supporting a concentration-dependent involvement of CaSR. Interestingly, cinacalcet reduced excitability primarily by lowering input resistance - a nonspecific effect not reported previously (Lindner *et al*., 2022), yet also limited the increase in AP number induced by low divalent cations.. Taken altogether, these results strongly support the conclusion that CaSR plays a critical role in the excitability change induced by variations in external divalent cation concentration.

The importance of CaSR in neuronal physiology aligns with its implication in multiple neurological disorders, including ischemic injury, Alzheimer’s disease, and epilepsy (Hannan *et al*., 2018). In particular, CaSR missense mutations is associated with idiopathic epilepsy syndrome (Kapoor *et al*., 2008) and low [Ca^2+^]_e_ can trigger epileptiform activity (Han *et al*., 2015). Our findings provide mechanistic insight into how CaSR dysregulation may contribute to pathological hyperexcitability.

### Ion channels involved in the increased excitability induced by low divalent cations

Pharmacological blockade of Nav1.2 channels revealed that this channel is a critical mediator of the excitability increase that accompanies a reduction in extracellular divalent cations. HWTx-IV significantly reduced the rheobase shift, and CRISPR-mediated knockdown of Nav1.1/Nav1.2 produced a comparable partial occlusion. In contrast, neither ATTx nor CRISPR-mediated Nav1.6 reduction altered excitability, suggesting that Nav1.6 does not contribute to threshold regulation under low divalent cation conditions. This discrepancy between the findings obtained in ATTx and in CRISPR-Nav1.6 may result from a homeostatic up-regulation of Nav1.2 when Nav1.6 is genetically reduced (Turrigiano & Nelson, 2004; O’Leary, 2018).

The genetic reduction of Nav1.1/2 has a significant impact on the augmentation in intrinsic excitability induced by the reduction of [Ca^2+^]_e_ and [Mg^2+^]_e_. Nav1.1 channels are mainly expressed in the axon of GABAergic interneurons and not in pyramidal cells (Lorincz & Nusser, 2008) while Nav1.2 channels are highly expressed in pyramidal cell axons (Hu *et al*., 2009). Therefore, the genetic deletion of Nav1.1/2 channels in our experiments is likely to exclusively target Nav1.2 channels in CA1 pyramidal neurons.

Previous work in dissociated neurons suggested that low [Ca^²⁺^]□ activates the leak Na⁺ channel NALCN, leading to the depolarization of the neuron (Xiong *et al*., 1997; Lu *et al*., 2010). In contrast, we observed no inward current during low divalent cation exposure; instead, a depolarizing holding current was required to maintain the membrane potential. This major difference is likely explained by the overrepresentation or deregulation of NALCN channels in dissociated neurons, possibly due to homeostatic changes. Furthermore, previous studies suggested the modulation of voltage-gated sodium channels in the enhanced excitability resulting from the deactivation of CaSR by low [Ca^2+^]_e_ (Mattheisen *et al*., 2018; Martiszus *et al*., 2021). However, in these studies the distinction between Nav1.2 and Nav1.6 had not been evaluated.

Kv channels also contributed to the AP threshold shift. Blocking Kv1 channels with DTx-I partially prevented the hyperpolarization of the AP threshold induced by low [Ca^²⁺^]□ and [Mg^²⁺^]□. While several potassium channels are inhibited by low [Ca^²⁺^]□ (Vassilev *et al*., 1997; Xu *et al*., 2005) Kv7 channels behave differently, being inhibited by CaSR activation or high [Ca^²⁺^]□ (Chuinsiri *et al*., 2024). Thus, different K⁺ channel families contribute in opposite ways to threshold regulation.

Together, these results identify CaSR as a promising target for modulating intrinsic excitability in conditions where extracellular Ca^²⁺^/Mg^²⁺^ homeostasis is perturbed. The ability of calcilytics to mimic and occlude the effects of low divalent cations suggests that intervention at the CaSR level could stabilize AP threshold dynamics in hyperexcitable states. Understanding how CaSR shapes the balance between Nav1.2-driven depolarization and Kv1-mediated repolarization may provide new therapeutic strategies for epilepsy and other disorders involving dysregulated neuronal excitability.

## Materials and methods

### Organotypic slice cultures of rat hippocampus

All experiments were carried out according to the European and Institutional guidelines for the care and use of laboratory animals and approved by the local health authority (D13055-08, Préfecture des Bouches-du-Rhône). Slices cultures were prepared as described previously (Debanne *et al*., 2008). In brief, young Wistar rats (P7–P10) were anesthetized with isoflurane and killed by decapitation, the brain was removed, and each hippocampus was dissected. Hippocampal slices (350 μm) were obtained using a Vibratome (Leica, VT1200S). They were placed on 20-mm latex membranes (Millicell) inserted into 35-mm Petri dishes containing 1 mL of culture medium and maintained for up to 18 d in an incubator at 35°C, 95% O_2_–5% CO_2_. The culture medium contained 25 ml MEM, 1.25 ml HBSS, 12.5 ml horse serum, 0.5 ml penicillin/streptomycin, 0.8 ml glucose (1 M), 0.1 ml ascorbic acid (1 mg/ml), 0.4 ml Hepes (1 M), 0.5 ml B27, and 8.95 ml sterile H_2_O.

### Electrophysiology

After 10-14 DIV, whole-cell recordings were obtained from CA1 pyramidal hippocampal neurons identified by the location of their soma, their morphology and their electrophysiological profile. Electroporated CA1 neurons were identified by GFP expression. Patch pipettes (7–9 MV) were filled with the internal solution containing (in mM): 120 K-gluconate, 20 KCl, 10 HEPES, 0 EGTA, 2 MgCl_2_, 2 Na_2_ATP, and 0.3 NaGTP (pH 7.4). All recordings were made at 29°C in a temperature-controlled recording chamber (Luigs & Neumann GmbH) with oxygenated ACSF containing (in mM): 125 NaCl, 26 NaHCO_3_, 3 CaCl_2_, 2.5 KCl, 2 MgCl_2_, 0.8 NaH_2_PO_4_, 0.6 Hepes and 10 D-glucose, equilibrated with 95% O_2_–5% CO_2_. Slice cultures were kept intact (i.e., without surgical removal of area CA3) and recordings from CA1 pyramidal neurons were made in the presence of intact inhibition (without PiTx) Neurons were recorded in current clamp with a Multiclamp 700B Amplifier (Molecular Devices). Excitability was measured by delivering a range of long (1 s) depolarizing current pulses (10– 290 pA, by increments of 10 pA) and counting the number of action potentials. In most experiments, the ratio of CaCl_2_ and MgCl_2_ concentration was maintained constant, and [Ca^2+^]_e_ was set to either 3 mM ([Mg^2+^]_e_ = 2 mM), 1.3 mM ([Mg^2+^]_e_ = 0.8 mM), or 0.6 mM ([Mg^2+^]_e_ = 0.4 mM).

Input-output curves were determined for each neuron, and three parameters were examined: the rheobase (the minimal current eliciting at least one action potential), the amplitude of the Action potential and the latency of the first spike (depolarizing time before the evoked spike under rheobase current eliciting only 1 spike). The current signals were low-pass filtered (10 kHz,), and acquisition was performed at 10 kHz with pClamp10 (Molecular Devices). Data were analyzed with ClampFit (Molecular Devices) and Igor (Wavemetrics). Spike thresholds were measured using phase plots (Fékété *et al*., 2021).

### CRISPR/Cas9

The primers used to design the specific gRNA targets were: Nav1.1, 1.2 forward: (5′ to 3′) CACC G tccactccccacacagcacg; Nav1.1, 1.2 reverse (3’ to 5’) AAAC cgtgctgtgtggggagtgga C; Nav 1.6 forward: (5′ to 3′) CACC G agtttgcctttcatctacg; Nav1.6 reverse (3’ to 5’) AAAC cgtagatgaaaggcaaact C; CaSR forward: (5′ to 3′) CACC G ctgctactccaaaatggat; CaSR reverse (3’ to 5’) AAAC atccattttggagtagcaag C. The gRNA sequences were ligated into pX458 to coexpress the human codon-optimized Cas9 as previously described (Incontro *et al*., 2014). We used pX458 expressing gRNA targeting Nav1.1, 1.2, pX458 expressing gRNA targeting Nav1.6, pX458 expressing gRNA targeting CaSR and FUGW expressing plasmid to aid identification of transfected cells. The human codon-optimized Cas9 and chimeric gRNA expression plasmid (pX458) as well as the lentiviral backbone plasmid (lentiCRISPR) both developed by the Zhang lab (Cong *et al*., 2013; Ran *et al*., 2013) were obtained from Addgene. To generate guide RNA (gRNA) plasmids, a pair of annealed oligos (20 bp) was ligated into the single gRNA scaffold of pX458 or lentiCRISPR. For all the experiments, we sub-cloned the two gRNAs including the following chimeric RNA into a pCAGGS-IRES-GFP plasmid, through PCR amplification and insertion into the plasmid using BstbI restriction sites. To enhance the identification of transfected neurons we co-expressed the pFUGW vector expressing only GFP with pCAGGS-IRES-GFP constructs.

### Single cell electroporation

Electroporation-mediated transfection (Rathenberg *et al*., 2003) was conducted in organotypic slice cultures of rat hippocampus at 4 d in vitro (DIV). Before electroporation, plasmid solutions were centrifuged at 10,000 g for 5 min to avoid obstruction of the micropipette. For single-cell electroporation (SCE), the microscope chamber consisted of a sterile 35-mm Petri dish. The plasmid DNA constructs were diluted to a final concentration of 33 ng/ul in the internal solution containing (in mM): 120 K-gluconate, 20 KCl, 10 HEPES, 0 EGTA, 2 MgCl_2_, 2 Na_2_ATP, and 0.3 NaGTP (pH 7.4). Micropipettes (7–10 MΩ) were filled with this DNA preparation after filtration with a sterile Acrodisc Syringe Filter (0.2 mm in pore diameter). During the SCE procedure, slice culture (DIV–DIV4) was positioned in the 35-mm Petri dish and covered with prewarmed and filtered external solution containing (in mM): 125 NaCl, 26 NaHCO_3_, 3 CaCl_2_, 2.5 KCl, 2 MgCl_2_, 0.8 NaH_2_PO_4_, 0.6 Hepes and 10 D-glucose, equilibrated with 95% O_2_–5% CO_2_. The ground electrode and the microelectrode were connected to an isolated voltage stimulator (Axoporator 800A, Molecular Devices). Under visual guidance, the micropipette was positioned by a three-axis micromanipulator near the cell body of selected CA1 neurons. Pressure was controlled to have a loose-seal between the micropipette and the plasma membrane. When the resistance monitored reached 25–30 MΩ, we induced a train of-12 V pulses during 500 ms (pulse width: 0.5 ms, frequency: 50 Hz). Each organotypic slice culture underwent SCE procedure for 5–10 selected neurons during a limited time of 30 min and was then back transferred to the incubator.

### Pharmacology

NPS 2143 hydrochloride (2-Chloro-6-[(2R)-3-[[1,1-dimethyl-2-(2-naphthalenyl)ethyl]amino-2-hydroxypropoxy]benzonitrile hydrochloride), Cinacalcet hydrochloride (N-[(1R)-1-(1-Naphthyl)ethyl]-3-[3-(trifluoromethyl)phenyl]propan-1-amine hydrochloride), 4,9-Anhydrotetrodotoxin (4S,5aS,6S,8R,9S,10S,11S,11aR,12R)-2-Amino-1,4,5a,6,8,9,10,11-octahydro-9-(hydroxymethyl)-6,10-epoxy-4,8,11a-metheno-11aH-oxocino[4,3-f][1,3,5]oxadiazepine-6,9,11-triol; ATTx) were purchased from Tocris. Huwentoxin IV (HWTx IV; C_174_H_277_N_51_O_52_S_6_) was purchased from Smartox Biotechnology. Kynurenate was obtained from Sigma-Aldrich and dendrotoxin-I (DTx-I) from Alomone Labs.

### Immunohistochemistry

Organotypic cultures from Wistar rats were processed as described previously (Extrémet *et al*., 2023). Briefly, slices were fixed in a solution containing 4% of paraformaldehyde in PBS for 15 min at 4°C, incubated in 50 mM NH_4_Cl in PBS for 15 min at room temperature (RT), and blocked overnight at 4°C in a solution containing 5% normal goat serum (NGS; Vector Laboratories) and 0.5% Triton X100 in PBS. After blocking, slices were incubated (24 h; 4°C) with primary antibodies in a solution containing 0.5% Triton X-100 and 2% NGS in PBS. The following antibody was used: mouse anti-CaSR (1:500, ThermoFisher, MA1-934, RRID: AB_325374). The slices were then washed four times for 20 min each time in PBS 0.5% Triton X-100 and then incubated with the appropriate secondary antibodies for 2 h at RT in a solution containing 0.5% Triton X-100 and 2%NGS in PBS. The following antibody was used: Alexa Fluor 488 goat anti-mouse (1:1000, ThermoFisher, A28175, RRID: AB_2536161). Subsequently, sections were washed three times for 20 min in PBS 0.5% Triton X-100. Nuclei were stained using DAPI at a final concentration of 1.5 mg/ml in PBS for 10 min and washed in PBS for 15 min. Slices were then mounted in Vectashield mounting medium (Vector Laboratories). Images were acquired as z-stacks with appropriate excitation and emission filters, 1 AU pinhole, and 25X oil immersion objective on a Zeiss LSM-780 confocal scanning microscope. Images were taken with 1.0 zoom, 1024*1024 pixels in z stacks with 1.0 μm steps. Laser power and gain settings were adjusted to prevent signal saturation.

### Western blots of cortical dissociated cultures

Mouse cortical neurons that contain ∼10% of glial lineages were obtained from embryonic brains using standard procedures. Briefly, timed-pregnant mice were sacrificed by cervical dislocation, and the embryos were extracted at embryonic day 16 (E16.5). Brains were quickly separated, cortices dissected, and meninges removed in a solution of cold Hank’s Balanced Salt Solution (HBSS) with 1% Penicillin-Streptomycin (P-S, Gibco) and then digested with 0.25% trypsin (Gibco) and DNAseI (Sigma-Aldrich) for 10 min at 37°C. The tissue was mechanically dissociated, centrifuged at 1000g for 3 min, and resuspended in CO_2_-equilibrated Neurobasal (Thermo Fisher Scientific) neuronal culture medium supplemented with 10% normal horse serum (NHS, Gibco), 1% Penicillin-Streptomycin, 0.5 mM L-glutamine, 1% P-S, and 5.8 μL NaHCO_3_/mL (Gibco). The cell suspension was pre-plated in an uncoated cell culture dish at 37 °C for 40 minutes to remove adherent glial cells. The pellet was resuspended in Neurobasal medium supplemented with 1% NHS, 1% P-S, 0.5 mM L-glutamine, 22 μM glutamic acid (Sigma-Aldrich), 2% B27 (Gibco), and 5.8 μL NaHCO_3_/mL, and plated at 50,000 cells per cm^2^ directly on tissue culture plates coated with poly-D-lysine (Sigma-Aldrich). After 24 hours, the medium was replaced with serum-free neuronal culture medium (1% P-S, 0.5 mM L-glutamine, 2% B27, 5.8 μL NaHCO_3_/mL).

Mouse cortical neurons were transfected 24 hours after plating using Polyplus Jet Optimus (VWR) at a DNA:lipid ratio of 1:1. Briefly, 500,000 cortical neurons seeded on PDL-coated 6-well plates were transfected with 1.3 µg of CaSR gRNA or empty vector, with an additional 100 ng of Turbo-GFP to monitor transfection efficiency. Neurons were transfected for 4 h and then a full media change was performed. Neurons were harvested 12 days after transfection, with half media change every 2 days.

Protein extracts were obtained from primary cultures at 12 DIV using RIPA buffer supplemented with protease and phosphatase inhibitors (Thermo Fisher Scientific, Halt protease and phosphatase inhibitor cocktail) and separated using SDS-polyacrylamide gel electrophoresis and proteins transferred to PVDF membranes. Blotting was performed using 5% non-fat milk for blocking and incubation with primary antibodies. Primary antibodies were incubated overnight at the following concentrations: b-Actin (1:6,000, Sigma-Aldrich #A2228) and CaSR (1:1,000, Thermo Fisher Scientific #PA585137). Secondary HRP-conjugated antibodies were incubated at room temperature for 45 minutes (1:5000, Thermo Fisher Scientific) and washed with TBS-T. Protein signals were detected by the ECL chemiluminescent system (Azure Biosystems). Densitometry analysis, standardized to b-Actin as a control for protein loading, was performed with Fiji software. Quantitation was performed from a minimum of two independent experiments with three technical replicates per condition.

### Statistics

Pooled data are presented as mean ± SEM and statistical analysis was performed using the Mann–Whitney *U*-test or Wilcoxon rank-signed test. Data were considered statistically significant for p < 0.05.

## Acknowledgments

We thank K. Milton and A. Venture for animal care and, N Boumedine-Guignon, L Fronzaroli-Molinieres, C Iborra-Bonnaure, S Brustlein and M Sangiardi for excellent technical assistance.

Author contributions: K.M. designed research, performed research, analyzed the data, built the figures and wrote the paper. M.A.S. performed research and analyzed the data. M.R. analyzed the data. D.D. designed research, analyzed the data, built the figures, provided funding and wrote the paper. S.I. designed and performed research, analyzed the data and wrote the paper.

## Data availability

All data needed to evaluate the conclusions in the paper are present in the manuscript and/or the Supplementary Materials.

---

**Figure S1.**
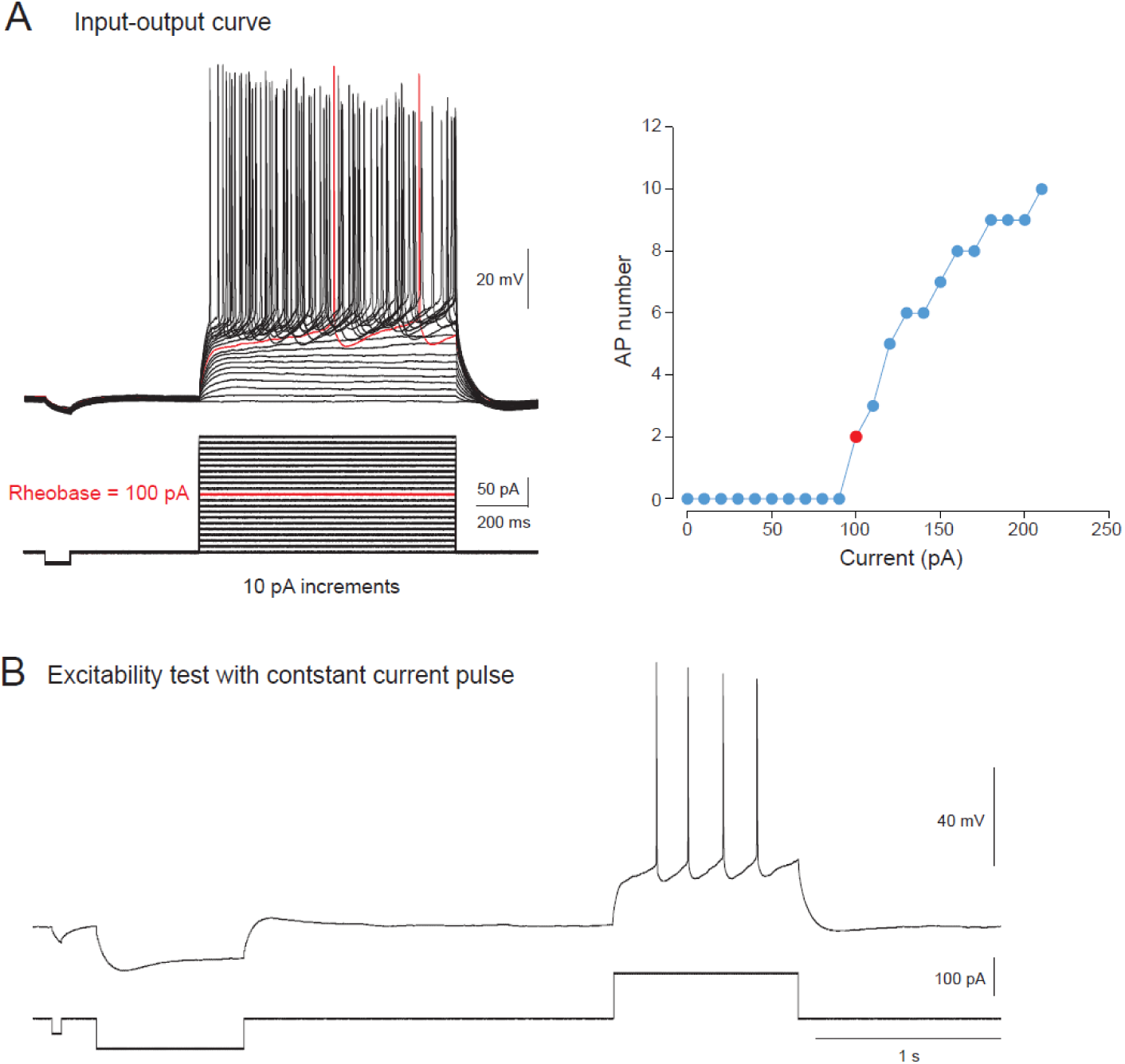
**Protocols of excitability exploration**. A. Input-output curve. Left, family of voltage traces in response to current pulse incremented by 10 pA. Right, input-output curve for the corresponding voltage traces. The rheobase is defined as the minimal current step for which at least 1 action potential is elicited. B. Test of excitability with constant current pulse used during the transitions from 3.0/1.3 mM calcium to 0.6 mM calcium. Red dot, rheobase.

**Figure S2.**
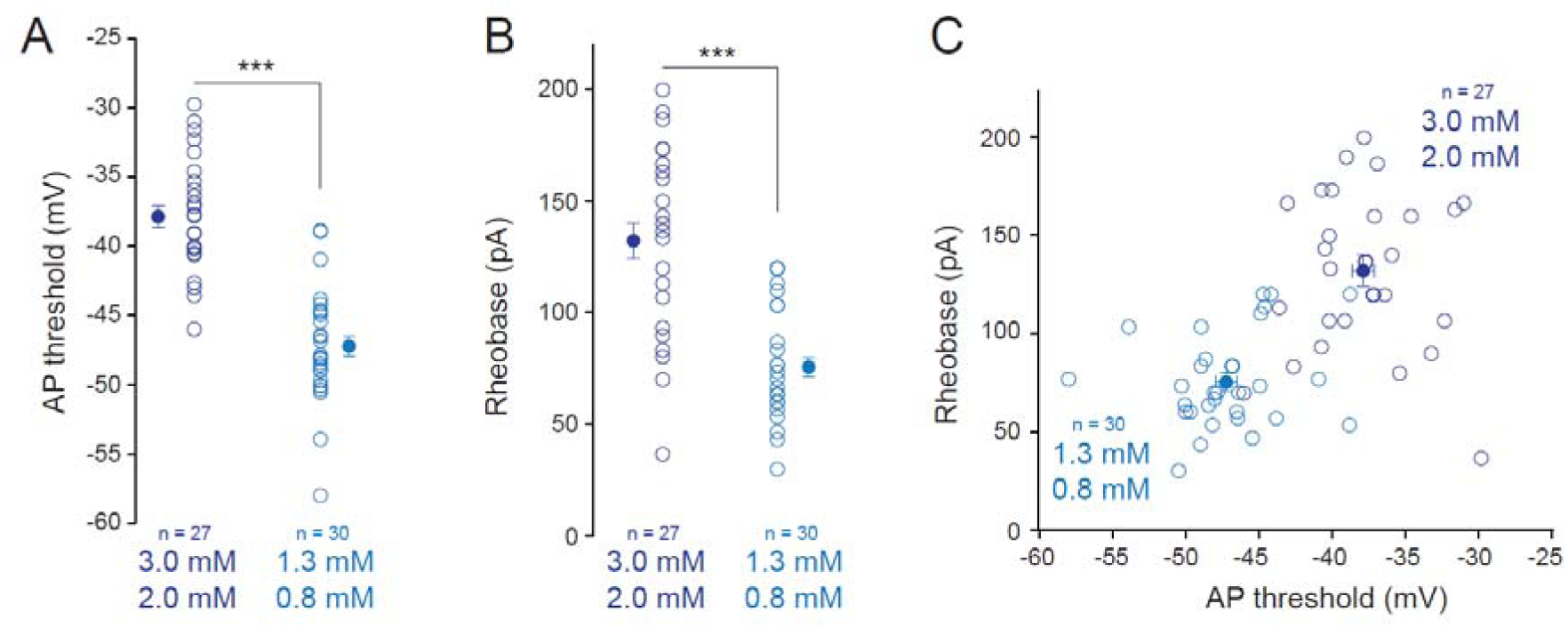
AP threshold and rheobase in high and physiological [Ca^2+^]_e_ and [Mg^2+^]_e_. A. Comparison of the AP threshold in high (3.0 mM / 2.0 mM) and physiological (1.3 mM / 0.8 mM) [Ca^2+^]_e_ and [Mg^2+^]_e_. ***, Mann-Whitney U-test, p<0.001). B. Comparison of the rheobase in high (3.0 mM / 2.0 mM) and physiological (1.3 mM / 0.8 mM) [Ca^2+^]_e_ and [Mg^2+^]_e_. ***, Mann-Whitney U-test, p<0.001). C. Plot of the rheobase vs. AP threshold for the data presented in A and B.

**Figure S3.**
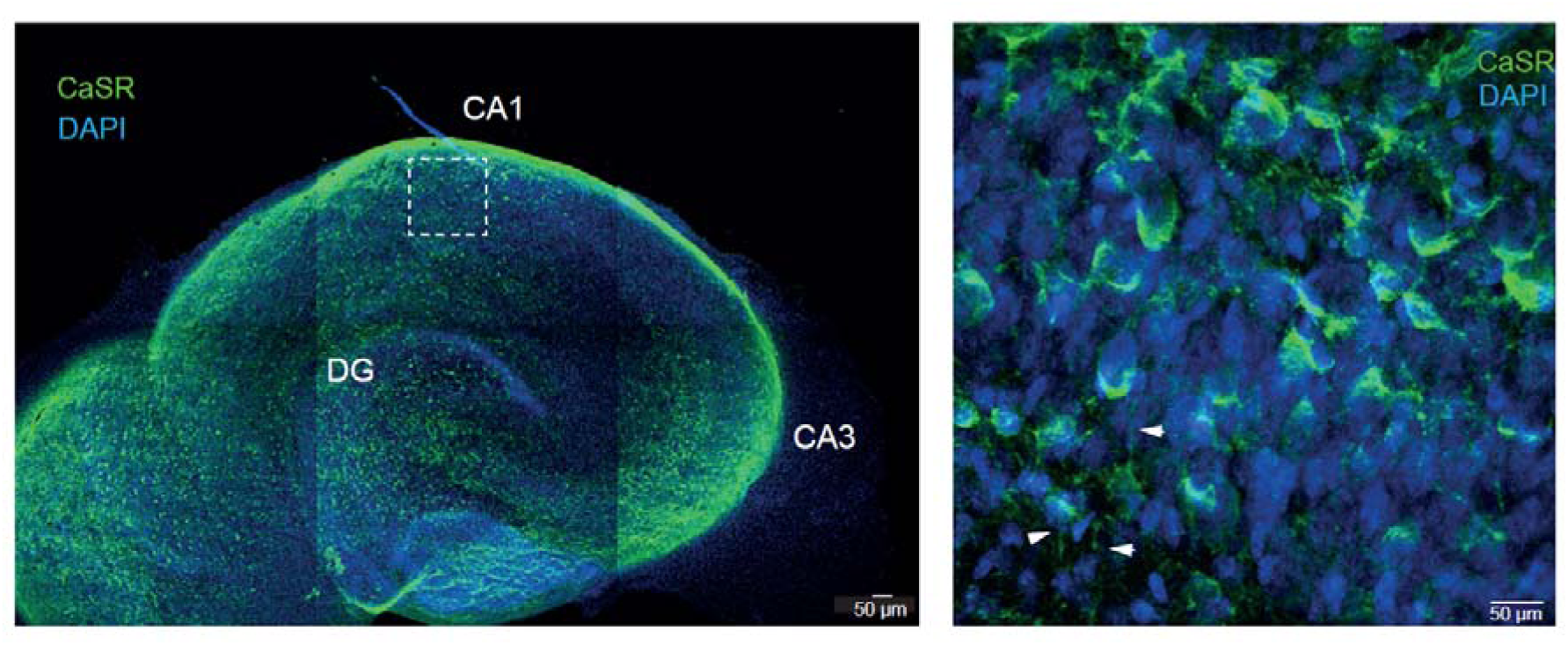
Immunostaining of CaSR in the area CA1 of hippocampal slice culture. Left, general view of the hippocampal slice culture stained with antibody against CaSR (green) and DAPI (blue). Right, magnification of the *stratum pyramidale* revealing the labelling in the cell bodies and in the apical dendrites (white arrow-heads).

**Figure S4.**
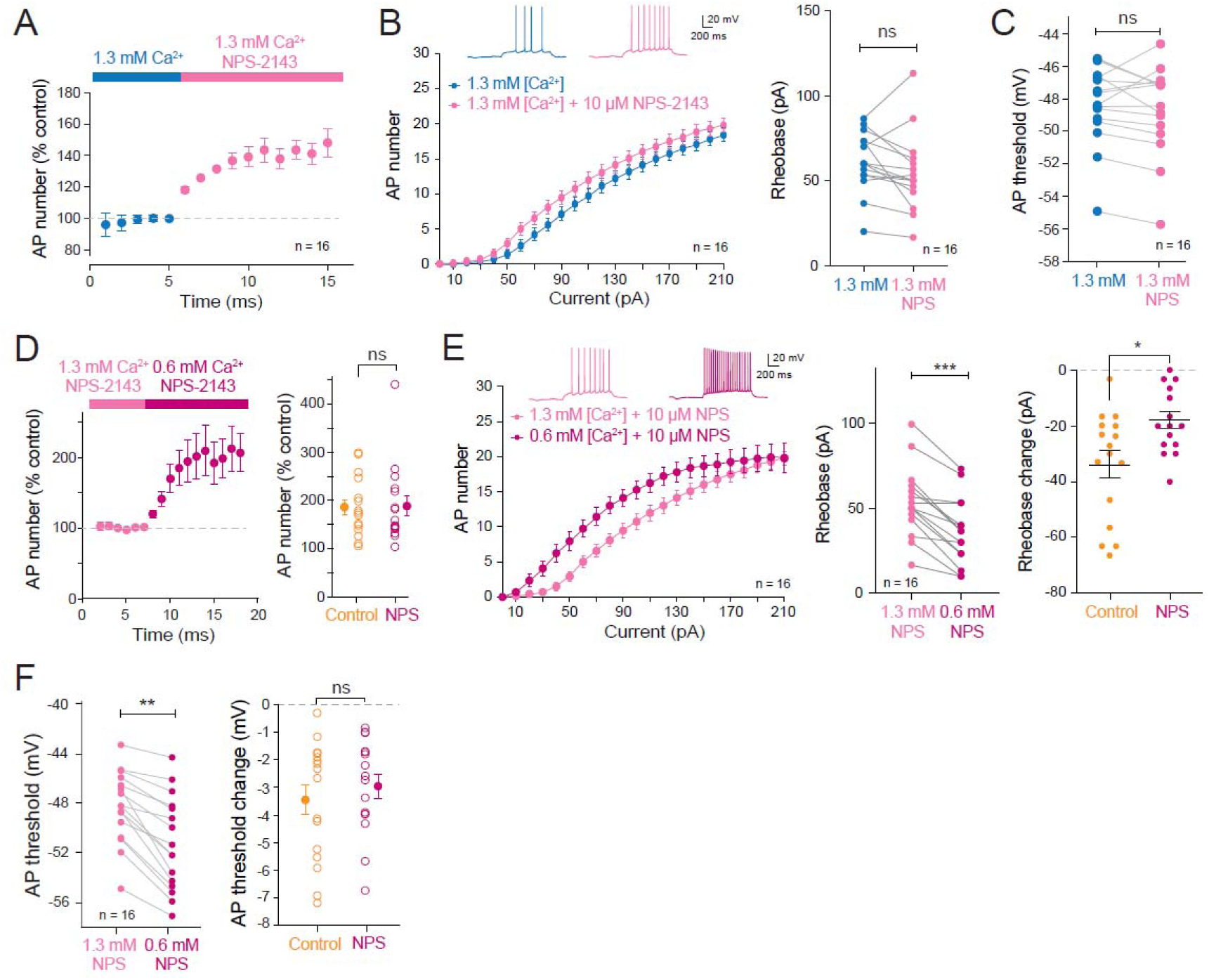
Effects of NPS-2143 in 1.3 mM Ca^2+^. A. Time-course of the increase in excitability caused by NPS-2143 in 1.3 mM Ca^2+^ and 0.8 mM Mg^2+^. B. Input-output curves in control (blue points) and in the presence of 10 µM NPS-2143. ns, Wilcoxon test, p>0.05. C. AP threshold analysis. ns, Wilcoxon test, p>0.05. D. Left, time course of the increase in excitability induced by low divalent cations. Right, comparison of AP number with the control. ns, Mann-Whitney U-test, p>0.1. E. Input-output curves in 1.3/0.8 mM (light pink) and in 0.6/0.4 mM (dark pink). Middle, Rheobase change. ***, Wilcoxon test, p<0.001. Right, comparison of the rheobase change with the control. *, Mann-Whitney U-test, p<0.05. F. AP threshold analysis. Left, hyperpolarization of the AP induced by low divalent ions. **, Wilcoxon test, p<0.01. Right, comparison of the AP threshold hyperpolarization with the control. ns, Mann-Whitney U-test, p>0.1.

**Figure S5.**
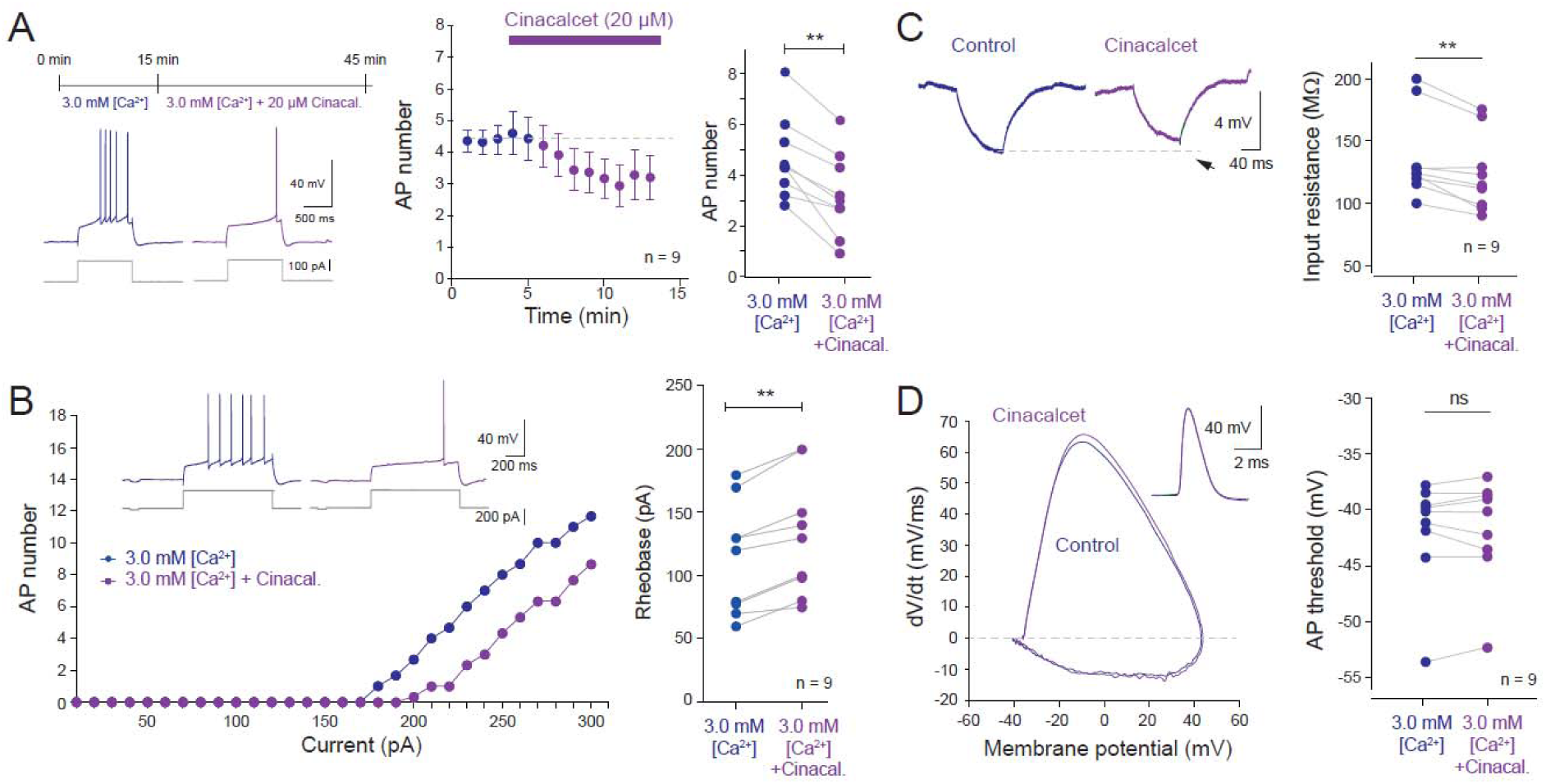
Cinacalcet reduces intrinsic excitability through reduced input resistance. A. Top left, protocol. Bottom left, example of firing inhibition induced by 20 µM cinacalcet. Middle, time-course of the reduction in neuronal excitability induced by cinacalcet. Right, statistical analysis. **, Wilcoxon test, p<0.01. B. Left, input-output curve shit induced by cinacalcet. Right, rheobase change. **, Wilcoxon test, p<0.01. C. Reduction of input resistance. Left, representative traces. Right, analysis. **, Wilcoxon test, p<0.01. D. AP threshold analysis. Left, phase plots in control and in cinacalcet. Right, AP threshold change. ns, Wilcoxon test, p>0.1.

**Figure S6.**
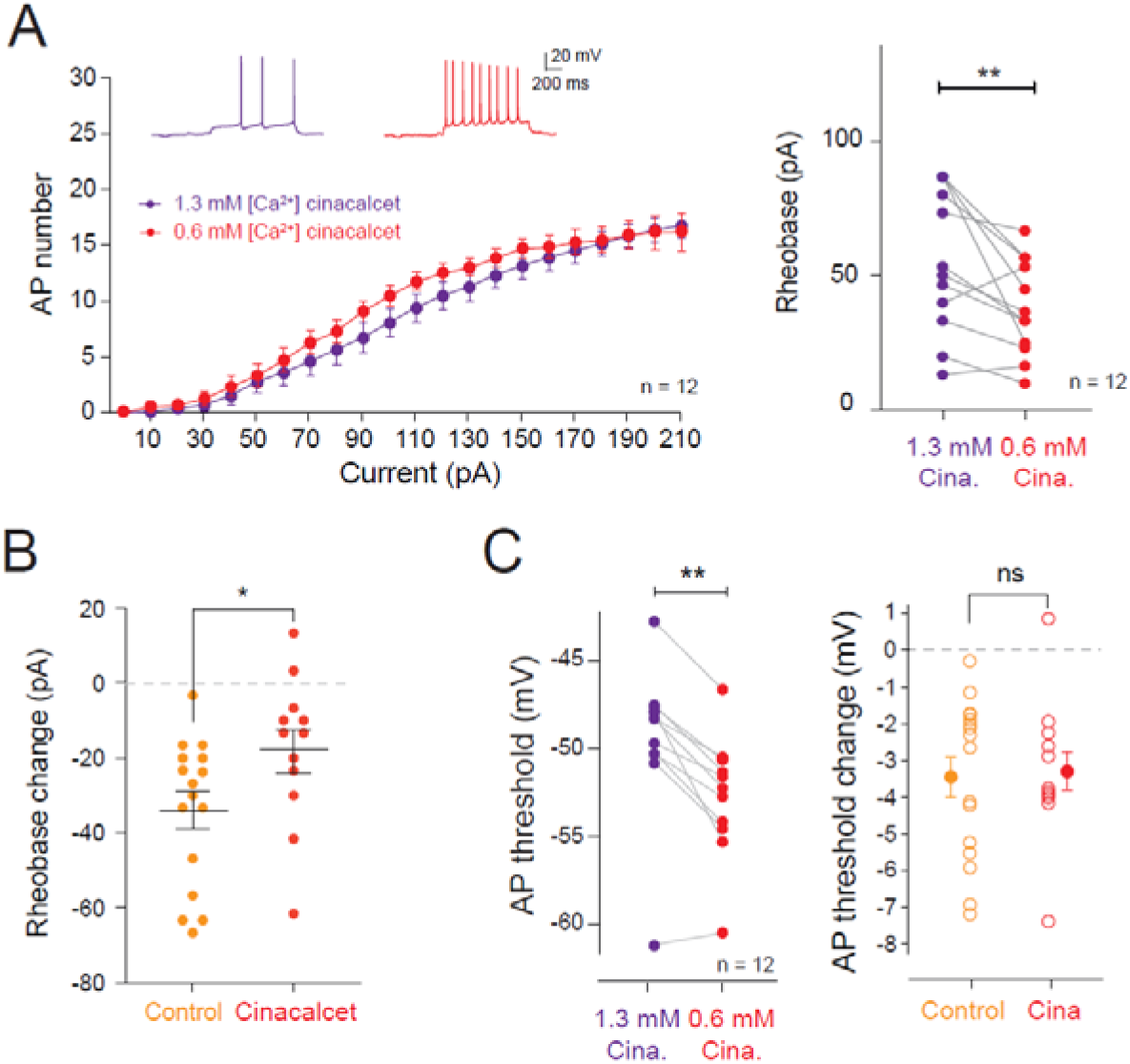
Cinacalcet partially occludes the rheobase reduction induced by the drop of divalent cations. A. Left, input-output curves in the presence of the calcimimetic cinacalcet. Right, reduction of the rheobase. **, Wilcoxon, p<0.01. B. Comparison of the rheobase reduction with the control. *, Mann-Whitney, p<0.05. C. Left, hyperpolarization of the AP threshold induced by low divalent cations in the presence of cinacalcet. **, Wilcoxon test, p<0.01. Right, comparison with the control. ns, Mann-Whitney U-test, p>0.1.

**Figure S7.**
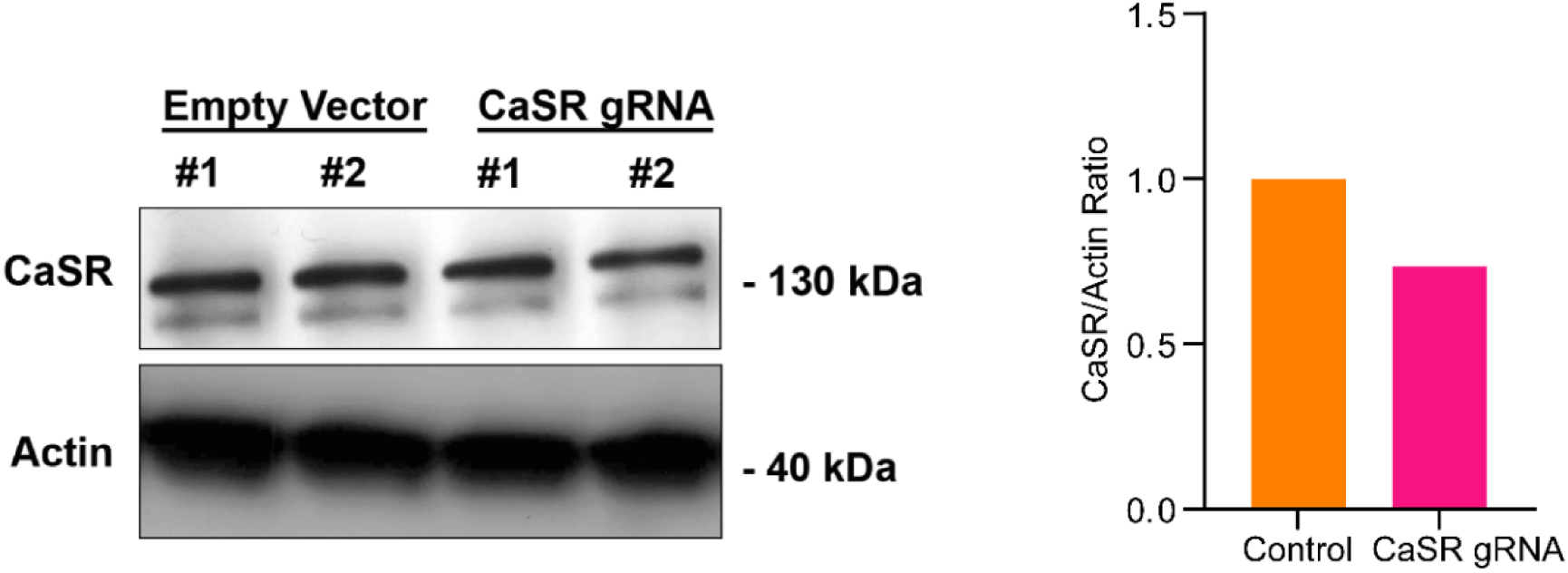
Western blot of CaSR in control and neurons transfected with the CRISPR CaSR. Top, CaSR signals measured in neurons transfected with an empty vector (left) and in neurons transfected with CaSR gRNA (right). Bottom, actin signals measured in the same samples and in a similar configuration. Right panel: quantification of the CaSR vs Actin ratio in control empty vector (1) and CRISPR/Cas9 CaSR transfected cells (0.7).

**Figure S8.**
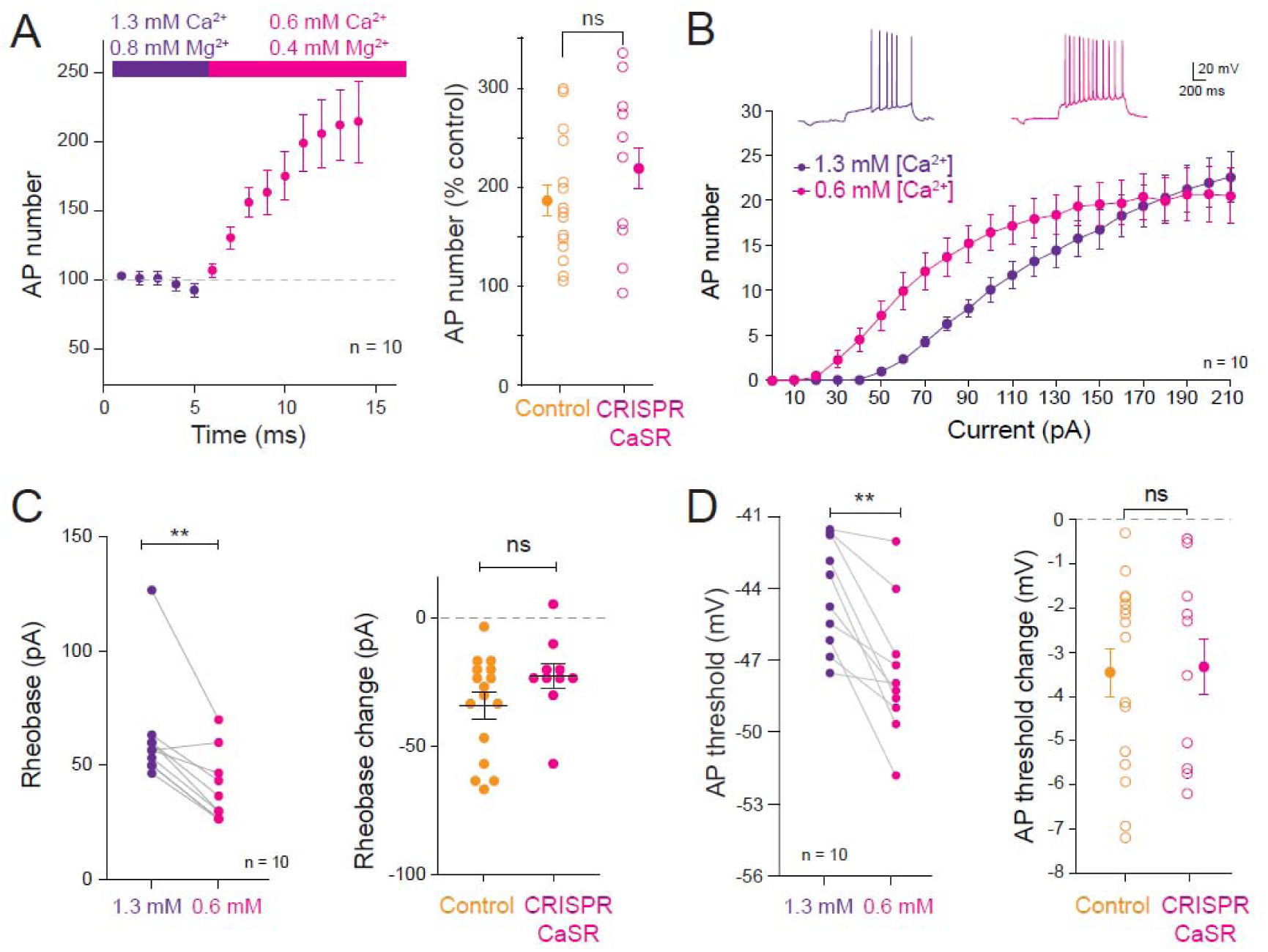
**Genetic impairment of CaSR does not occlude excitability enhancement induced by low divalent cations in physiological conditions**. A. Time-course of the excitability enhancement induced by low divalent cations (left). Right, comparison of the AP number in control and in CRISPR CaSR neurons. ns, Wilcoxon, p>0.1. B. Input-output curves in physiological conditions and in the presence of low divalent cations. C. Left, rheobase changes induced by low calcium and magnesium concentration. **, Wilcoxon test, p<0.01. Right, comparison with the control. ns, Mann-Whitney U-test, p>0.05. D. Left, AP threshold changes. **, Wilcoxon test, p<0.01. Right, Comparison with the control. ns, Mann-Whitney U-test, p>0.1.

**Figure S9.**
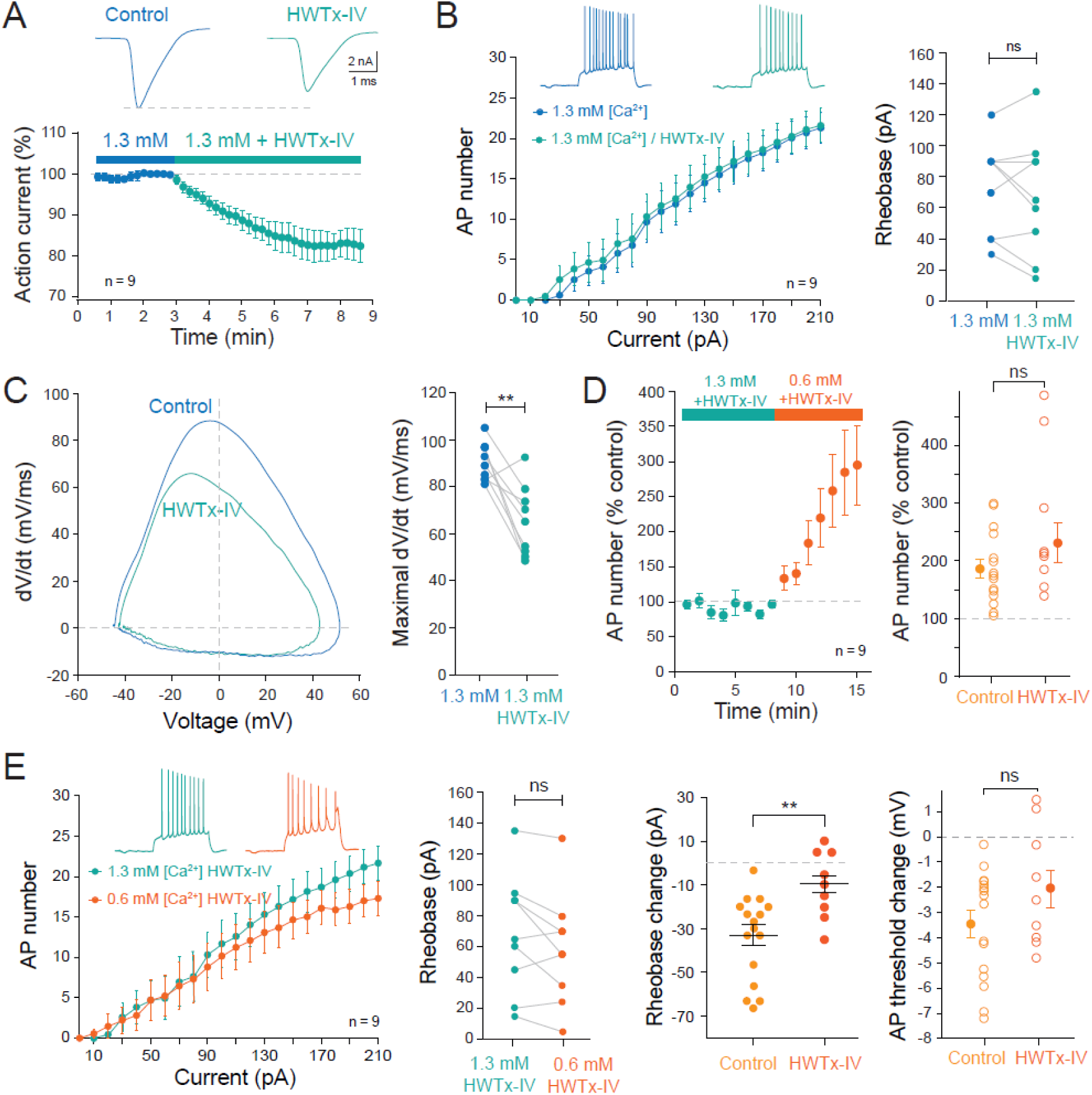
HWTx partially occludes the increase in excitability in physiological conditions. A. Reduction of the Nav current by 300 nM of HWTx-IV. Top, representative currents evoked by a depolarization from-80 to 0 mV in a CA1 pyramidal neuron. Bottom, time-course of the reduction. B. HWTx-IV does not alter the input output curve. Left, input-output curves. Right rheobase. ns, Wilcoxon test, p>0.1. C. AP phase-plot (left) and maximal dV/dt changes (right) induced by low divalent cations in the presence of HWTx-IV. **, Wilcoxon test, p<0.01. D. Time course of the enhanced excitability induced by low divalent cations (left) and comparison with control (right). ns, Mann-Whitney U-test, p>0.1. E. Left, input-output curves in high and low divalent cations. Middle left, changes in the rheobase. ns, Wilcoxon test, p>0.05. Middle right, comparison of the rheobase changes in control and in HWTx. Wilcoxon test, p<0.01. Right, comparison of the AP threshold hyperpolarization in HWTx and in control. ns, Mann-Whitney U-test, p>0.05.

**Figure S10.**
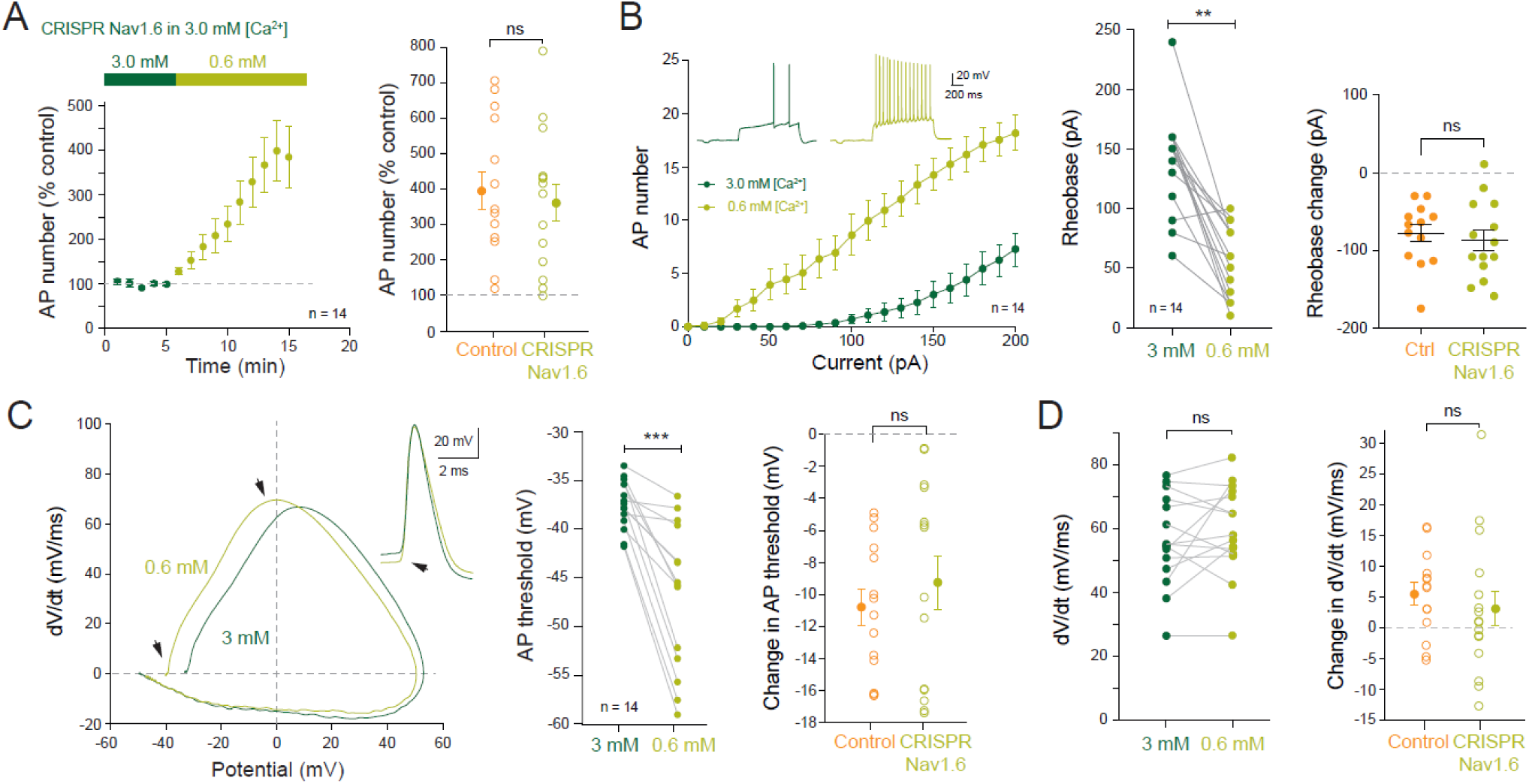
Genetic impairment of Nav1.6 does not alter the elevation in intrinsic excitability. A. Time-course of the excitability enhancement induced by low divalent cations in CRISPR Nav1.6 neurons (left). Right, comparison of the AP number in control and in CRISPR Nav1.6 neurons. ns, Wilcoxon, p>0.1. B. Input-output curves in high and in low divalent cations left). Middle, rheobase changes induced by low calcium and magnesium concentration. **, Wilcoxon test, p<0.01. Right, comparison with the control. ns, Mann-Whitney U-test, p>0.1. C. AP threshold changes. Left, AP phase-plots in the presence of high and low divalent cations. Middle, AP threshold change. ***, Wilcoxon test, p<0.001. Right, Comparison with the control. ns, Mann-Whitney U-test, p>0.1. D. Maximal dV/dt analysis. Left, changes in maximal dV/dt. ns, Wilcoxon test, p>0.1. Right, comparison of the changes in dV/dt with the control. ns, Mann-Whitney U-test, p>0.1.

**Figure S11.**
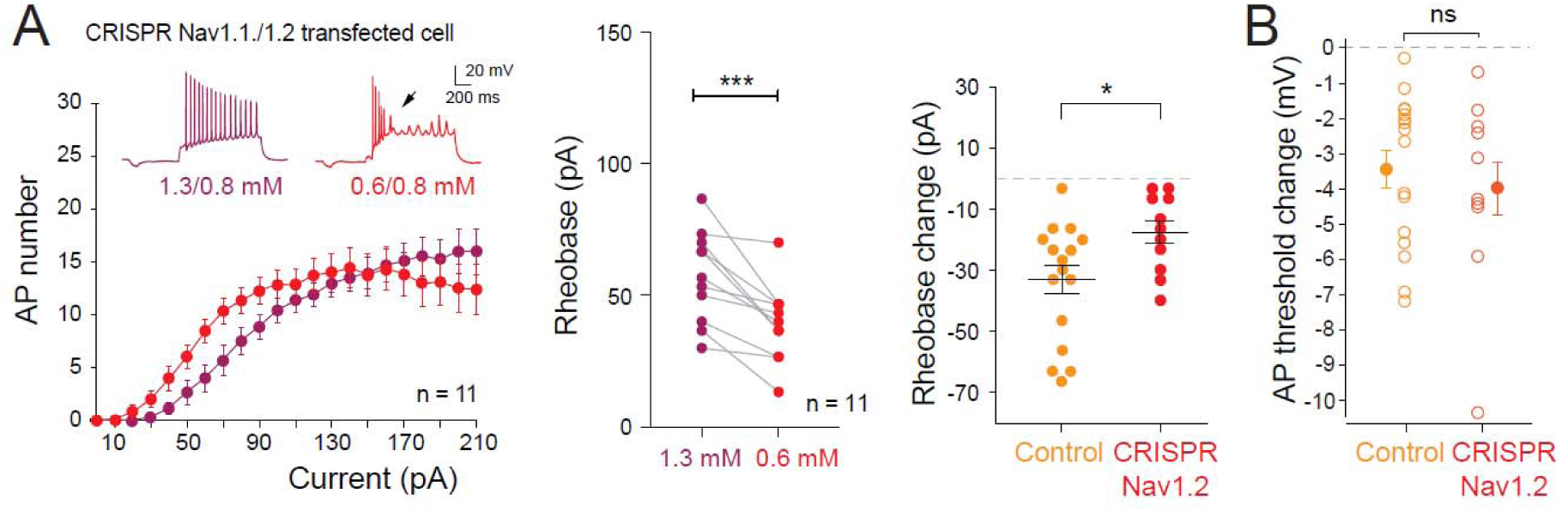
Genetic impairment of Nav1.1/Nav1.2 partially occludes the enhanced excitability caused by [Ca^2+^]_e_ and [Mg^2+^]_e_ drop. A. Left, input-output curves in 1.3/0.8 mM (purple) and in 0.6/0.8 mM (red) in CRISPR Nav1.1/Nav1.2 transfected neurons. ***, Wilcoxon test, p<0.001. Note the failures of AP after the initial discharge (arrow). Middle, rheobase modification. Right, comparison of the change in rheobase in control and in CRISPR Nav1.2. *, Mann-Whitney U test, p<0.05. B. Comparison of the AP threshold change in control and in CRISPR Nav1.2. ns, Mann-Whitney U test, p>0.1.

